# APOE2, E3 and E4 differentially modulate cellular homeostasis, cholesterol metabolism and inflammatory response in isogenic iPSC-derived astrocytes

**DOI:** 10.1101/2021.07.17.452644

**Authors:** Sherida M. de Leeuw, Aron W. T. Kirschner, Karina Lindner, Ruslan Rust, Witold E. Wolski, Anne-Claude Gavin, Roger M. Nitsch, Christian Tackenberg

## Abstract

Apolipoprotein E (APOE) is the principal lipid carrier in the CNS and mainly expressed by astrocytes. The three different *APOE* alleles (E2, E3, and E4) impose differential risk to Alzheimer’s disease (AD); E2 is protective, E3 is defined as average risk, while E4 is the major genetic risk factor for sporadic AD. Despite recent advances, the fundamental role of different *APOE* alleles in brain homeostasis is still poorly understood. To uncover the functional role of APOE in human astrocytes, we differentiated human *APOE*-isogenic iPSCs (E4, E3, E2 and APOE-knockout (KO)) to functional astrocytes (hereafter “iAstrocytes”), with a resting, non-proliferating phenotype. Functional assays indicated that polymorphisms in APOE (APOE4>E3>E2=KO) reduced iAstrocyte metabolic and clearance functions including glutamate uptake and receptor-mediated uptake of β-amyloid aggregates. We performed unlabelled mass spectrometry-based proteomic analysis of iAstrocytes at baseline and after activation with interleukin-1β (IL-1β) showing a reduction of cholesterol and lipid metabolic and biosynthetic pathways, and an increase of immunoregulatory pathways at baseline (E4>E3>E2). Cholesterol efflux and biosynthesis were reduced in E4 iAstrocytes, and subcellular localization of cholesterol in lysosomes was increased. In *APOE*-KO iAstrocytes, APOE-independent mechanisms showed to be proficient in mediating cholesterol biosynthesis and efflux. Proteomic analysis of IL-1β-treated iAstrocytes showed an increase of cholesterol/lipid metabolism and biosynthesis as well as inflammatory pathways. Furthermore, cholesterol efflux, which was reduced in *APOE4* iAstrocytes at baseline, was alleviated in activated E4 iAstrocytes. Inflammatory cytokine release was exacerbated upon IL-1β treatment in E4 iAstrocytes (E4>E3>E2>KO), in line with the proteomic data. Taken together, we show that APOE plays a major role in several physiological and metabolic processes in human astrocytes with *APOE4* pushing iAstrocytes to a disease-relevant phenotype, causing dysregulated cholesterol/lipid homeostasis, increased inflammatory signalling and reduced β-amyloid uptake while *APOE2* iAstrocytes show opposing effects.

Our study provides a new reference for AD-relevant proteomic and metabolic changes, mediated by the three main APOE isoforms in human astrocytes.

## Introduction

Apolipoprotein E (APOE) is part of a family of lipoprotein chaperones, binding lipoprotein complexes and facilitating their uptake through low-density lipoprotein receptors (LDLRs), or lipoprotein receptor 1 (LRP1), providing lipids and cholesterol to the cell (Kockx et al., 2018). In turn, APOE aids in cholesterol and lipid efflux from the cell by associating with lipoprotein particles upon formation (Mahley, 2016). APOE is expressed throughout the body, produced mainly by the liver and the brain. In the brain APOE is one of the most abundant lipoprotein chaperones, mainly expressed by astrocytes, and in low levels by microglia.

APOE mainly exists in 3 allelic variants, APOE2, E3, and E4, which differ in a single amino acid change at position 112 or 158; APOE2 (Cys112, Cys158), APOE3 (Cys112, Arg158), and APOE4 (Arg112, Arg158) (Liu et al., 2013). Positions 112 and 158 are located in helix 3 and 4, respectively, where they lie adjacent to the receptor binding region of the APOE protein. The Cys112 residue in APOE2 reduces its receptor binding capacity, congruent with the increased levels of lipoproteins in plasma, leading to hyperlipidemia III (Fernandez et al., 2019). Secondly, Arg158 in APOE4 causes an ionic interaction between Arg61 and Cys255 on the lipid binding domain, inferring a structural change, leading to altered lipid binding and decreased capacity to load cholesterol.

Carrying the *APOE4* allele is the major genetic risk factor to develop sporadic Alzheimer’s disease (AD), where one allele confers a 3-fold increase and two alleles a 12-fold increase in risk. On the other hand, carrying an *APOE2* allele is protective for AD, however, its biological role is heavily understudied and not yet understood (Michaelson, 2014). AD is the most common age-related neurodegenerative disorder and is characterized by the presence of β-amyloid (Aβ) plaques and neurofibrillary tangles. The accumulation and aggregation of Aβ, caused by reduced clearance mechanisms, plays a significant role in the progression of AD pathogenesis (Winblad et al., 2016). Early AD pathology is signified by a complex cellular phase, presenting a myriad of dysfunctions such as synapse loss, elevated immune response, defective clearance by the endolysosomal system, ER stress, and an increase in cholesteryl esters, cholesterol, and decreased levels of glycerophospholipids (De Strooper and Karran, 2016; Henstridge et al., 2019; Kao et al., 2020; Tackenberg et al., 2009). It has been hypothesized that these dysfunctions may be preceded by loss of homeostatic control of astrocytes (Rodríguez-Arellano et al., 2016). Considering that APOE is primarily expressed in astrocytes, a key role is implied for astrocytic APOE in the initial phase of AD pathophysiology.

We aimed to understand the fundamental role of the three different APOE isoforms (APOE2, E3, E4), in human iPSC-derived astrocytes. After successfully deriving non-proliferative astrocytes at a resting state, we show that *APOE2, E3, E4* and APOE-KO iAstrocytes show distinct phenotypes in homeostatic functions, cholesterol and lipid metabolism, lysosomal function, and inflammatory regulation, in addition to the interaction of these mechanisms. While several cellular functions, such as glutamate uptake, seem to display a clear allele-dependent decrease, the interplay of cholesterol metabolism, subcellular localization and immune regulation indicate not a mere functional decrease in *APOE*4 iAstrocytes: *APOE2, APOE3* and *APOE4* may use different mechanisms altogether. Additionally, we show that iAstrocytes utilize a primary APOE-dependent cholesterol metabolic mechanism, as well as highly efficient APOE-independent secondary mechanisms.

Collectively our data provides novel insights into the role of the three different APOE alleles on fundamental human astrocyte biology, indicating key roles in astrocyte homeostasis, cholesterol and lipid function, lysosomal function, inflammatory response as well as the interaction of inflammatory regulation and cholesterol homeostasis.

## Results

### Robust differentiation of quiescent iAstrocytes from iPSC-derived neural progenitor cells

There is currently an abundance of differentiation protocols for iPSC-astrocytes. However, the majority are laborious; require tedious cocktails of small molecules, sorting methods presuming previous expertise, and use fetal calf serum (FCS), resulting in activated astrocytes at baseline (Li et al., 2018; Lundin et al., 2018; Santos et al., 2017; Zhang et al., 2016). By simply applying ScienCell Primary Astrocyte Medium (AM) to neural progenitor cells (NPCs), TCW and colleagues (TCW et al., 2017) generated functional iPSC-astrocytes in 30 days, overcoming the complicated methods currently used, albeit still using FCS. We have therefore extended this method by withdrawing FCS and adding cytosine arabinoside (AraC) to the medium at day 30, for 7 days. Followed by another 7 days of AM without FCS and AraC, taking a total of 44 days; generating mature and quiescent iAstrocytes (**Fig1A, F**).

**Figure 1.**
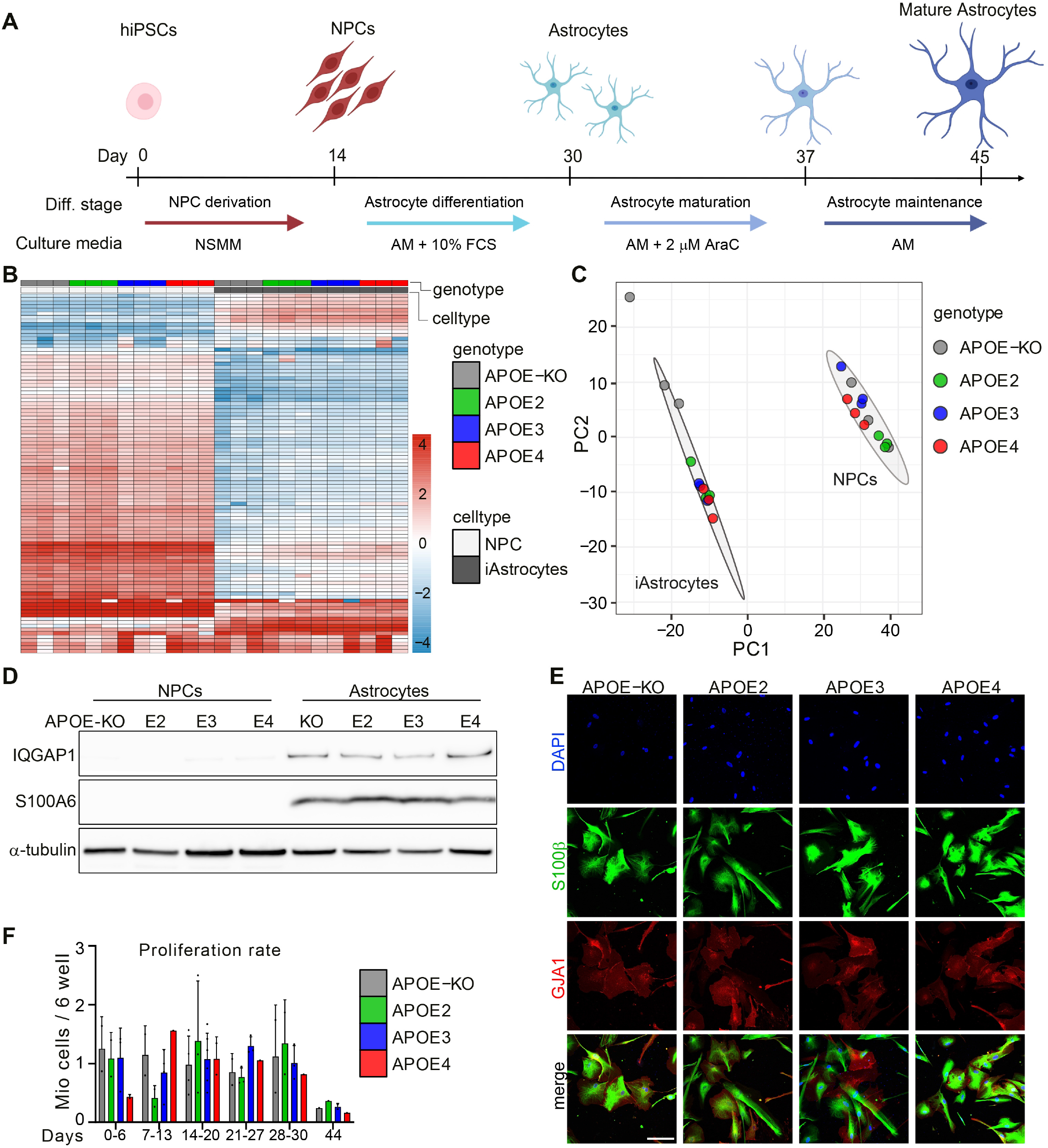
APOE isogenic iPSCs are differentiated to iAstrocytes. A: Schematic representation of the iAstrocyte differentiation protocol. B: Heatmap of top 100 differentially expressed proteins between NPCs and iAstrocytes selected by fold-change. C: PCA of proteomic data from NPCs and iAstrocytes of all genotypes. Three biological replicates were analyzed. D: Western blot of astrocyte marker proteins IQGAP1 and S100A6 in APOE-isogenic NPCs and Astrocytes. E: lmmunocytochemistry of astrocyte markers S100β and GJA1 in APOE-isogenic iAstrocytes at day 44. Scale bar: 150μm. F: Proliferation rate of APOE-isogenic iAstrocytes at different time points in culture. NSMM: Neural Stem cell Maintenance Medium, AM: astrocyte medium

Isogenic *APOE* iPSCs were obtained from EBiSC, and show mono-allelic expression of *APOE*, or APOE deletion (Schmid et al., 2019; 2020). Unlabelled mass spectrometry (MS)-based proteomic analysis of iAstrocytes and NPCs from 4 different genotypes (*APOE2, APOE3, APOE4* and *APOE-KO*; (**FigS1A**)), resulted in a total of 3765 detected proteins, of which the majority, 2432 proteins, were differentially expressed in iAstrocytes compared to NPCs (FDR < 0.1), from which the 100 most differentially expressed proteins were visualized (**Fig1B**). Principal component analysis showed separate clustering of NPCs and iAstrocytes, indicating all *APOE* isogenic lines were successfully differentiated from NPCs to iAstrocytes (**Fig1C**). It should be noted that *APOE-KO* iAstrocytes show more genotype variation than E2, E3 and E4 iAstrocytes, and we observed a substantially lower number of peptides and proteins in mass-spectrometry analysis. Since this would bias the data normalization for the *APOE*-KO samples, we omitted them from the quantitative proteomic analysis. Among the astrocyte markers significantly upregulated in iAstrocytes compared to NPCs in the proteomic data were calcium binding protein S100A6 and Rho-ATPase IQGAP1, which were confirmed with Western blot showing robust expression in iAstrocytes, and absence in NPCs (**Fig1D**). Immunocytochemical analysis of mature astrocyte markers, calcium binding protein S100β and gap junction protein GJA1, showed all cells being positive at day 44 (**Fig1E**), as well as for intermediate filament protein GFAP (**FigS1B**). Compared to iAstrocytes at day 44, cells on day 30 displayed a different morphology (**FigS1C**); the latter were smaller and expressed most GJA1 in the cytosol, while GJA1 was prominently expressed on the cell surface on day 44, in addition to the significant increase in cell volume demonstrated in the S100β staining. Altogether we show successful differentiation of NPCs to quiescent iAstrocytes, carrying *APOE2, APOE3, APOE4* or *APOE-KO* alleles, showing a mature phenotype.

### Homeostatic functions show APOE allele-dependent decline in iAstrocytes

To investigate *APOE-*related phenotypes in isogenic iAstrocytes, proteomic analysis of *APOE2, APOE3, APOE4* iAstrocytes was performed showing most differently regulated proteins (fold change above 1 or below -1) between *APOE2* and *APOE4* (173), less between *APOE2* and *APOE3* (130) and least between *APOE4* and *APOE3* (72) (**Fig2A**). We analyzed APOE levels in iAstrocytes which were regulated in isoform-dependent manner (E2>E3>E4), and absent in *APOE-KO* iAstrocytes (**Fig2B, C**). In line with the extracellular lipoprotein-binding function of APOE, we observed a larger amount of APOE protein in the secreted fraction, where the difference between genotypes was higher than in the lysate. APOE is known to affect many functional aspects of astrocytes, including, but not limited to, homeostatic support such as uptake of glutamate as well as uptake and degradation of Aβ, as shown in mouse models (Verghese et al., 2013; Zhong et al., 2008).

**Figure 2.**
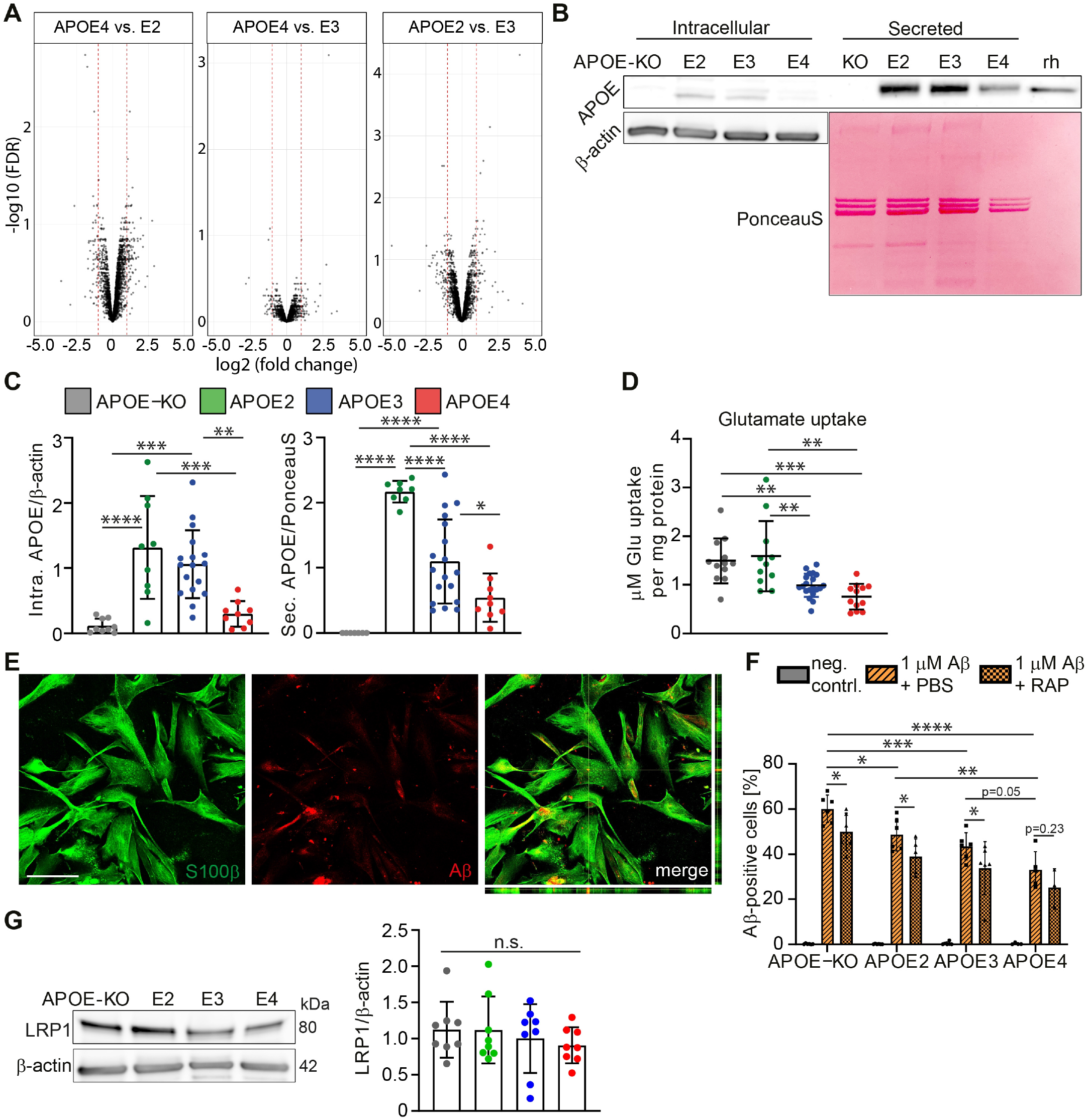
APOE isogenic iAstrocytes show allele-dependent decline in homeostatic functions. A: Volcano plots of proteomic analysis comparing APOE2 vs APOE3, APOE4 vs APOE3 and APOE4 vs APOE2 iAstrocytes. Log2 of fold change is plotted against -log10 of FDR (false discovery rate). Red lines indicate a fold change of 1. B: Western blot of APOE in lysate (intracellular) and supernatant (secreted) of iAstrocytes. Intracellular APOE was normalized to β-actin, secreted APOE to the PonceauS signal of the respective lane. Recombinant human (rh) APOE served as control. C: Quantification of APOE western blot bands from lysate (Intra. APOE) and supernatant (Sec. APOE). D: Glutamate uptake assay showing the amount of glutamate taken up by APOE-isogenic iAstrocytes within 1hr. Data was normalized to total protein content of the respective cell lysate. E: Representative confocal images of stained iAstrocytes (S100β), treated with 1μM Aβ42. Scale bar: 150 μm. F: Flow cytometry-based Aβ42 uptake assay showing percentage A positive cells of live cells. Cells were treated with 1μM pre-aggregated Aβ42-hilyte488 or pre-treated for 24hrs with 0.2μM of LRP1 antagonist RAP. G: Western blot and quantification of LRP1 in lysates from iAstrocytes. Data represent mean± SD(*: p < 0.05; **: p < 0.01; ***: p < 0.001; ****: p < 0.0001, one-way ANOVA with post hoc Tukey’s multiple comparisons test (C, D, G), two-way ANOVA with post hoc Holm-Šídák’s multiple comparisons test (F)).

Uptake of glutamate in iAstrocytes demonstrated an allele-dependent effect with *APOE4* iAstrocytes taking up the lowest amount of glutamate while *APOE2 and APOE-KO* took up the highest amounts (**Fig2D**). Reduced glutamate uptake was not caused by the lower APOE levels in *APOE4* iAstrocytes, as the *APOE-KO* iAstrocytes showed similar glutamate uptake capacity as *APOE2*. As a second functional measure, we determined the ability of iAstrocytes to take up Aβ (**Fig2E**). A flow cytometry-based Aβ42 uptake assay, using pre-aggregated Hilyte488-tagged Aβ42, showed an allele-dependent trend in uptake capacity (E2>E3>E4), although differences between *APOE3* and *APOE4* did not reach significance (p=0.054) (**Fig2F**). Interestingly, *APOE-KO* cells showed the highest Aβ42 uptake capacity. Aβ42 is mostly taken up through receptor-dependent endocytosis, such as LRP1. To determine if Aβ uptake in iAstrocytes is mediated by LRP1-dependent endocytosis, we treated cells with a low concentration of LRP1 antagonist RAP in order to induce mild reduction in Aβ uptake resulting in a measurable difference between genotypes. Blocking LRP1 caused a significant decrease in Aβ uptake in *APOE-KO*, E2 and E3, but not in E4 iAstrocytes (p=0.23). Quantification of LRP1 protein levels showed no significant difference, albeit a minor decrease of average LRP1 levels in *APOE4* cells (**Fig2G**). These results demonstrate an isoform-dependent regulation of APOE protein levels, as was previously shown for E3 and E4 *in vitro*, or in patient CSF (Lin et al., 2018; Mahley, 2016). We further show an allele-dependent effect on iAstrocyte homeostatic functions (KO=E2>E3>E4), where *APOE2* closely resembled *APOE-KO* iAstrocytes.

### Cholesterol and lipid metabolism show APOE allele-dependent regulation

Gene Set Enrichment Analysis (GSEA) of proteomic data showed significant downregulation of pathways involved in lipid biosynthesis/metabolism and sterol biosynthesis/metabolism in *APOE*4 compared to *APOE*3 or *APOE*2 iAstrocytes, while these were upregulated in *APOE*2 compared to *APOE*3 (**Fig3A**). One of the proteins significantly regulated was FDFT1 (squalene synthase), the first committed enzyme in the *de novo* biosynthesis pathway of cholesterol, or mevalonate pathway (Do et al., 2009). A second protein of interest was ABCA1, ATP-ase binding cassette 1, essential for lipid and cholesterol efflux, as well as for APOE lipidation (Oram, 2003). Western blot analysis confirmed that these proteins were allele-dependently regulated (KO=E2>E3>E4) (**Fig3B, C**). To investigate the functional consequences of these observations, cholesterol, cholesteryl esters (CE) and phosphatidylethanolamines (PE) were quantified using high-performance thin-layer chromatography (HPTLC) from cell pellet and supernatant. In line with the proteomic data, cellular cholesterol was decreased in *APOE*4 compared to *APOE*3 and *APOE*2 iAstrocytes, while showing similar levels as APOE-KO (E2=E3>E4=KO) (**FigS2A**), whereas cellular CEs were only decreased in *APOE4* compared to E3. Cellular PE showed higher levels in E2 and E4 and TAG displayed an isoform-dependent increase (E4>E3>E2=KO). Interestingly, secreted cholesterol was significantly decreased only in *APOE*4 iAstrocytes (**Fig3D**). CEs and PEs did not show a genotype-dependent difference in cellular versus secreted lipids. The reduced secretion of cholesterol by the *APOE*4 iAstrocytes could be explained by a difference in subcellular localization. We therefore visualized non-membrane bound cholesterol by Methyl-β-Cyclodextrin (MβCD) treatment and subsequent Fillipin III staining, as well as additional Dextran-Alexa555 treatment, to determine lysosomal localization of cholesterol. We found that non-membrane bound cholesterol was increased in *APOE*4 iAstrocytes (E4>E3=E2=KO) (**Fig3E**). Further co-localization of Fillipin III with Dextran-Alexa555 was highest in *APOE4* iAstrocytes, indicating higher cholesterol levels in lysosomes of *APOE*4 iAstrocytes (E4>E2) (**Fig3F**). These results indicate that while there is a decrease of total cholesterol in *APOE*4 iAstrocytes, the fraction of non-membrane bound and lysosomal, cholesterol is higher compared to other genotypes. A reduced fraction of cholesterol is thus available for transport in concordance with a decrease of cholesterol efflux machinery ABCA1, in combination with a decrease of *de novo* synthesized cholesterol due to the lack of FDFT1. Opposed to *APOE*-KO which showed decreased level of cholesterol compared to E2 and E3, but not a significantly different fraction of non-membrane bound cholesterol.

**Figure 3.**
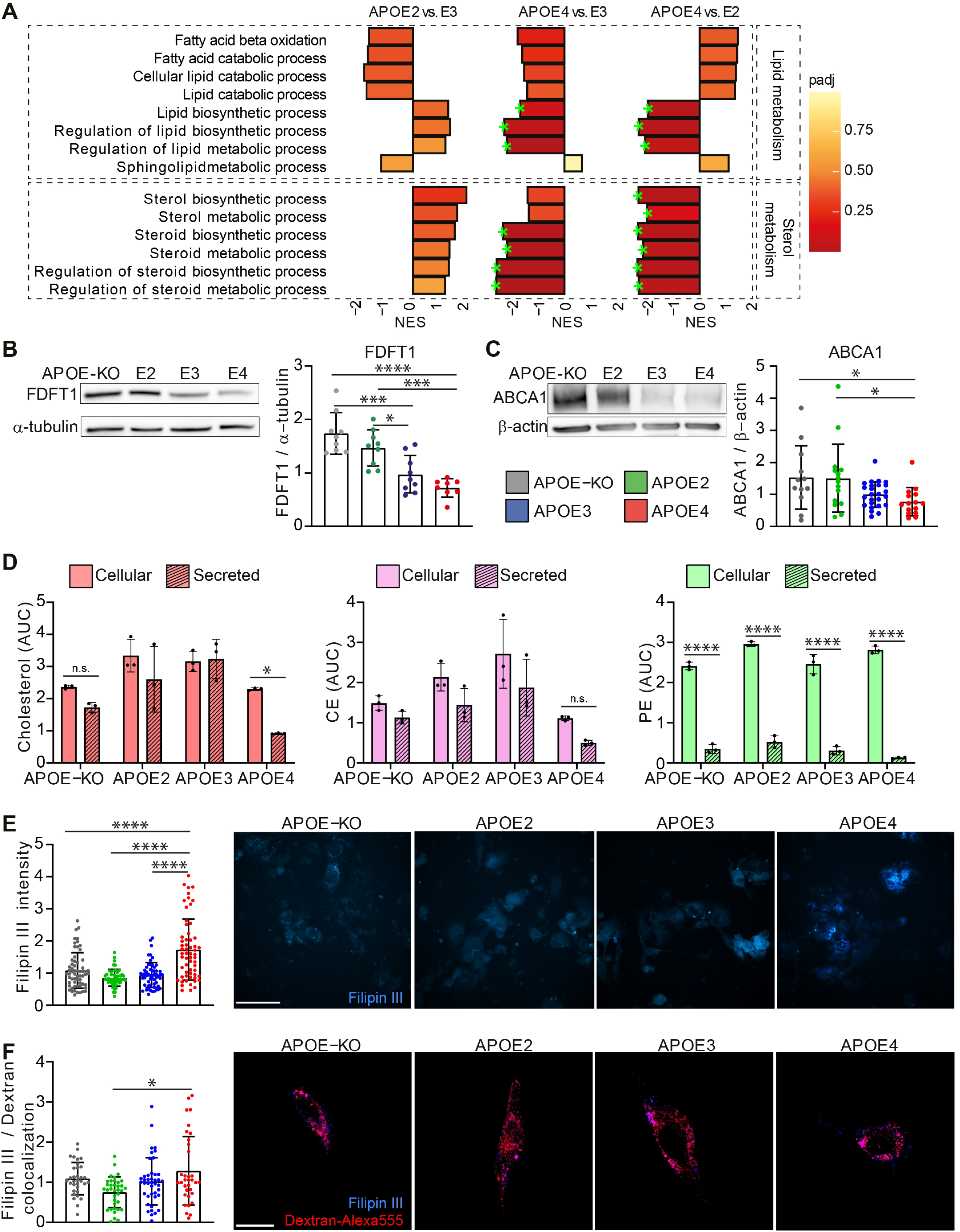
APOE isogenic iAstrocytes show allele-dependent regulation of cholesterol and lipid metabolism. A: GSEA Normalized Enrichment scores (NES) of protein lists ranked according to the t-statistic obtained for the contrast APOE2 vs APOE3, APOE4 vs APOE3 and APOE4 vs APOE2 iAstrocytes. NES is plotted on the x-axis, with color-coaded bars for the individual GO terms, according to adjusted p-value (padj). We annotated gene sets with padj < 0.2 (green stars). B: Western blot and quantification of FDFT1 in lysates from iAstrocytes, normalized to a-tubulin. C: Western blot and quantification of ABCA1 in lysates from iAstrocytes, normalized to β-actin. **D:** Cellular and secreted cholesterol, cholesteryl ester (CE) and phosphatydilethanolamine (PE) quantified with HPTLC, normalized to cellular phosphatidylcholine content. **E:** Bar graphs of Filipin III intensity quantification, normalized to *APOE3* intensity, with representative example images. Scale bar: 150µm. **F:** Bar graphs showing quantification of Filipin III and Dextran-Alexa555 colocalization in iAstrocytes. Representative images of Filipin III (blue) and Alexa555 (red) overlay are displayed. Scale bar: 50µm. Data represent mean ± SD (*: p < 0.05; ***: p < 0.001; ****: p < 0.0001, Kruskall-Wallis test (C, F), One-way ANOVA with *post hoc* Tukey’s multiple comparisons test (B, E) or two-way ANOVA with *post hoc* Holm-Šídák’s multiple comparisons test (D)).

### Lysosomal but not endosomal function is affected by APOE genotype

To further investigate possible defects in the endolysosomal system affecting degradation of extracellular waste such as Aβ, or cholesterol trafficking, relevant pathways in our proteomic data were analysed. Proton and cation transmembrane transport were highest in *APOE*4 iAstrocytes, and lowest in E2 (**Fig4A**). V-ATPase-regulated proton transport into the lysosomal lumen is crucial to maintaining the acidic pH, and defective acidification has been linked to AD or Parkinson’s disease (Colacurcio & Nixon, 2016). An increase in proton and cation transmembrane transport might indicate a compensatory mechanism for underlying dysfunction. In addition to these pathways, multiple ion homeostasis regulatory pathways were enriched in *APOE*4 but did not reach significance. Lysosomal compartments contain a plethora of ion transporters and channels to maintain their function (Xiong & Zhu, 2016). Positive regulation of ion transport and ion homeostasis pathways were enriched in *APOE*3 compared to *APOE*2, other ion regulatory pathways were not significantly regulated. In addition, *APOE*2 iAstrocytes showed moderate decrease of ion transporter, homeostatic or regulatory pathways.

**Figure 4.**
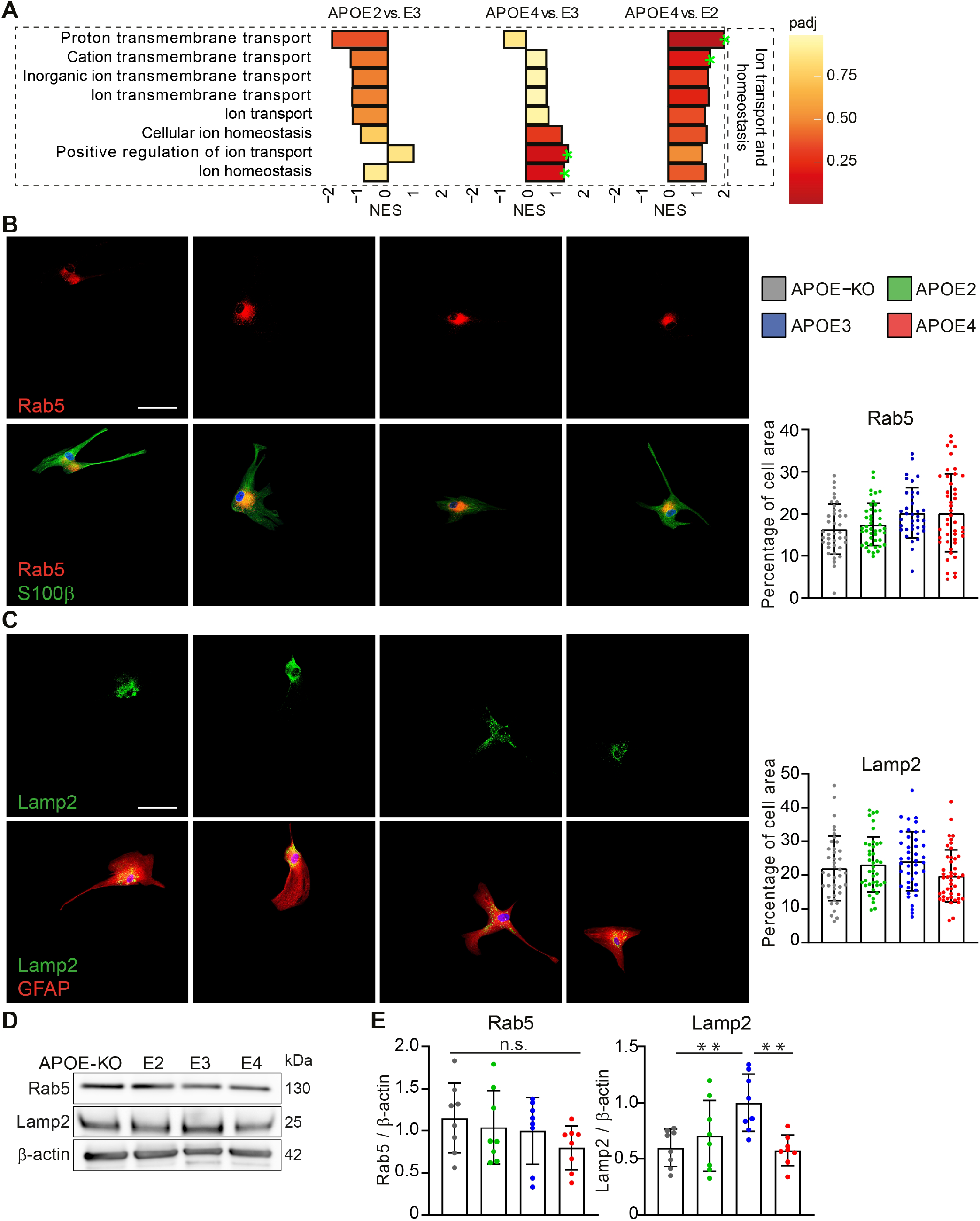
Subtle changes in lysosomal marker LAMP2 according to APOE genotype. A. GSEA NES of protein lists ranked according to the t-statistic obtained for the contrast APOE2 vs APOE3, APOE4 vs APOE3 and APOE4 vs APOE2 iAstrocytes. NES is plotted on the x-axis, with color-coded bars according to adjusted p-value (Padj) Padj ≤ 0.2 was considered significant, annotated with green stars. B. lmmunocytochemical stainings of RABS (red, upper panel) and merged with S100β (green, lower panel). The signal is quantified and calculated as percentage of cell area. C. lmmunocytochemical stainings of LAMP2 (green, upper panel) and merged with GFAP (red, lower panel). The signal is quantified and calculated as percentage of cell area. Scale bar: 50μm. D. Western Blot of LAMP2 and RABS from lysate of iAstrocytes. Scale bar: 50μm. E. Western blot quantification of RAB5 or LAMP2 normalized to β-actin. Data represent mean± SD(**: p < 0.01, ordinary one-way ANOVA with post hoc Holm-Holm-Šídák’s multiple comparisons test).

We further assessed the levels of lysosomal and endosomal markers LAMP2 and RAB5. Endosomal compartments labelled with RAB5 did not show a significant difference in percentage of cell area between genotypes, neither did RAB5 protein levels (**Fig4B, D, E**). Immunofluorescent detection of lysosomes by LAMP2 (**Fig4C**) or Dextran-Alexa555 (**FigS3**) showed no difference between genotypes. However, protein levels of LAMP2 detected by Western blot were significantly increased in *APOE*3 compared to *APOE*2, *APOE*4, *APOE*-KO (E3>E2=E4=KO) (**Fig4D, E**). These results show no major abnormalities in endosomal and lysosomal compartment morphology, while LAMP2 protein levels are affected by *APOE* genotype, possibly indicating the initiation of a pathological mechanism together with the insistence of a compensatory mechanism detected in proteomic pathway analysis.

### APOE genotype-dependent upregulation of inflammatory pathways and exacerbated cytokine release in IL-1β treated iAstrocytes

A hallmark of AD pathology is an increased inflammatory microenvironment (Heneka et al., 2015). Reactive microglia secrete cytokines, such as IL-1β or TNF-α, to induce a cellular inflammatory cascade, in turn affect and activate astrocytes. To assess the role of *APOE* genotype on the inflammatory state of astrocytes, we treated iAstrocytes with IL-1β and performed proteomic analysis. At baseline, pathways involved in inflammatory regulation were highest in APOE4 compared to E3 and E2 and lower in E2 compared to E3 (E4>E3>E2). (**Fig5A**).When stimulated with IL-1β, the majority of these pathways were downregulated in E2, but showed enrichment in E3 and E4 (**Fig5B**). On the contrary, regulation of exocytosis and NF-κB signalling pathways are significantly downregulated in *APOE4*, but not in *E2* or *E3*. GSEA showed an allele-dependent effect in differentially regulated pathways (E4>E3>E2): In *APOE4* and *APOE3* iAstrocytes inflammatory pathways were predominantly upregulated after IL-1β treatment, while the opposite was found in *APOE2* cells. Interestingly, the cell aging pathway was significantly upregulated in E4, enriched, but not significantly, in E3, and even less in E2. Lastly, cell-cell junction was significantly downregulated in E4 but only slightly in E2 and E3, indicating that an inflammatory microenvironment affects astrocytes, especially E4, on a fundamental level, from cell aging to cellular communication. Interestingly, *APOE4* iAstrocytes showed the most differentially expressed pathways (both up and down regulated), more than *APOE3*, and *APOE2* iAstrocytes showed almost no significantly regulated pathways, indicating that *APOE4* iAstrocytes are most susceptible to pro-inflammatory stimuli (**Fig5C**).

**Figure 5.**
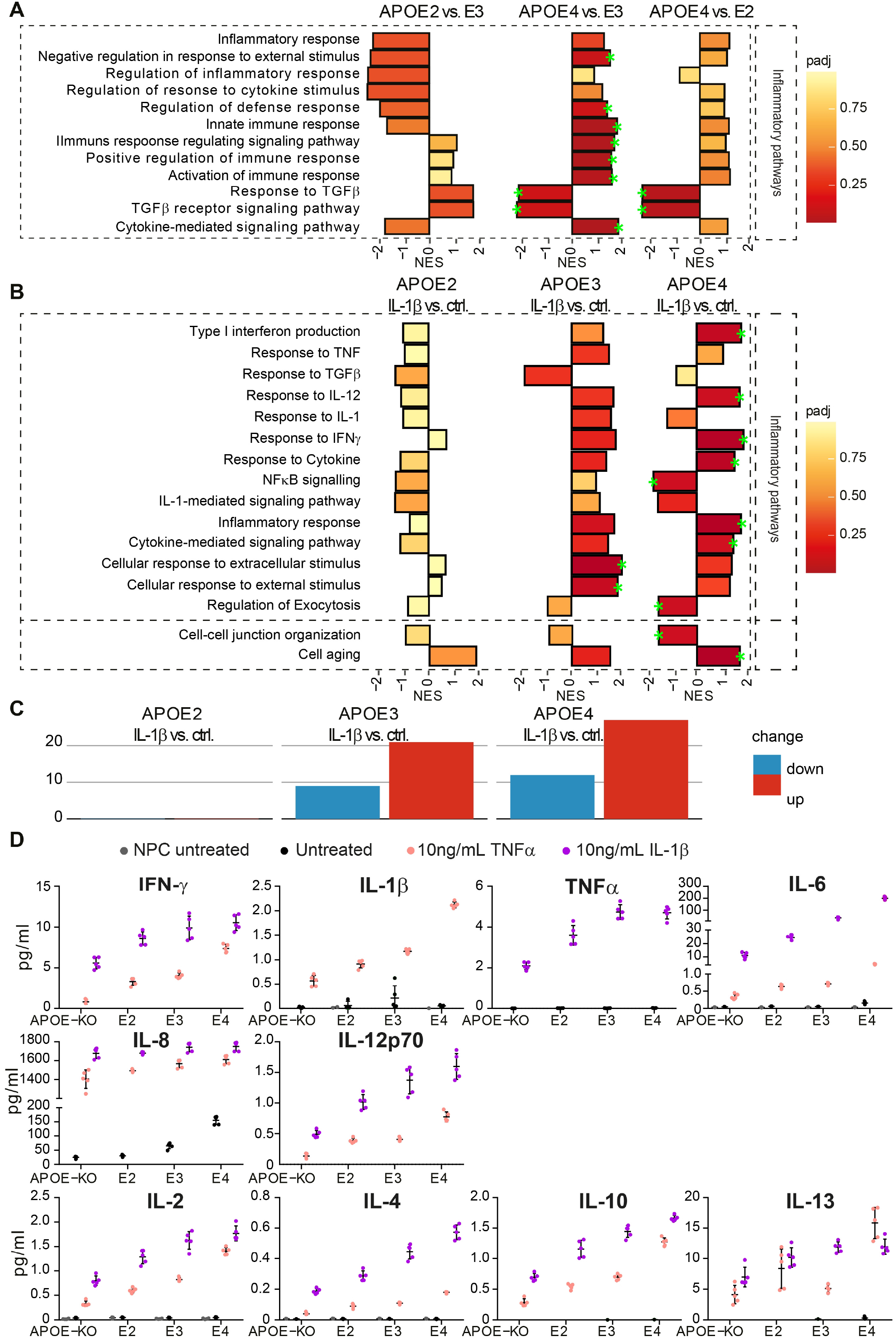
*APOE* genotype dependent effects on inflammatory pathway expression and cytokine release. **A**. GSEA Normalized Enrichment scores (NES) of protein lists ranked according to the t-statistic obtained for the contrast *APOE2* vs *APOE4, APOE4* vs *APOE3* and *APOE4* vs *APOE2* iAstrocytes respectively. **B**. GSEA of differential protein expression of baseline vs IL-1β treated iAstrocytes (*APOE2*, E3 and E4 respectively). NES is plotted on the x-axis, with color-coaded bars according to adjusted p-value (Padj). We annotated gene sets with padj < 0.2 (green stars). **C**. Number of differentially expressed biological pathways determined by GSEA, with a padj < 0.5. **D**. Levels of inflammatory cytokines secreted by NPCs at baseline (grey), iAstrocytes at baseline (black), 10ng/mL IL-1β-treated iAstrocytes (coral) and 10ng/mL TNF-α-treated iAstrocytes (lilac)., measured with MSD. Statistical analysis of MSD is depicted in **TableS1**.

Release of cytokines upon IL-1β or TNF-α treatment was measured using MSD, showing a strong allele-dependent release in all cytokines (E4>E3>E2>KO), except for IL-8 (**Fig5D, TableS1)**. This cytokine already showed detectable release at baseline, following a similar trend (E4>E3>E2=KO). Interestingly, IL-6 was increased upon stimulation, but also showed detectable levels at baseline in *APOE4* iAstrocytes. We then assessed whether IL-1β treatment also affected homeostatic functions by measuring Aβ42 uptake after IL-1β treatment of the iAstrocytes, but besides the previously shown allele-dependent degree uptake, no significant effect of the treatment was observed (**FigS4**). These data suggest that *APOE4* iAstrocytes have an inflammatory phenotype already at baseline which is exacerbated upon activation, while *APOE2* cells display a rather anti-inflammatory or neutral phenotype, and *APOE3* represents an intermediate state.

### IL-1β treatment affects cholesterol and CE regulation

Lastly, we assessed a potential interaction of IL-1β treatment and cholesterol, fatty acid and lipid metabolism/biosynthesis. These pathways were mostly downregulated in *APOE4* versus *APOE2* and *APOE3* at baseline **(Fig3A)**, but after IL-1β stimulation they were enriched in *APOE4*, and downregulated in *APOE2*, whereas *APOE3* showed differential regulation of these pathways (**Fig6A**). Lipid and cholesterol content was assessed after IL-1β treatment using HPTLC. In contrast to baseline, secreted versus cellular cholesterol was not different between genotypes anymore (**Fig6B**). Interestingly, IL-1β treatment induced a significant increase in cellular (**FigS2B**) but not in secreted cholesterol (**FigS2C**), only in *APOE3* iAstrocytes, indicating IL-1β treatment induces synthesis or uptake of cholesterol. Similar to baseline, IL-1β treatment showed highly significant secreted versus cellular PEs in all genotypes (**Fig6B**). To investigate whether IL1-β treatment affected non-membrane bound cholesterol, iAstrocytes were treated with IL-1β prior to MβCD treatment and Fillipin III staining. Opposed to the increase in non-membrane bound cholesterol at baseline, IL-1β treatment completely abolished the allele-dependent effect: levels of non-membrane bound cholesterol were similar in IL-1β treated iAstrocytes in all *APOE* genotypes (**Fig6C**). Altogether, these data indicate a role of the inflammatory microenvironment on cholesterol regulation with *APOE2* cells being unaffected, *APOE3* cells increasing their cholesterol content, and *APOE4* cells showing altered cholesterol regulation upon IL-1β treatment.

**Figure 6.**
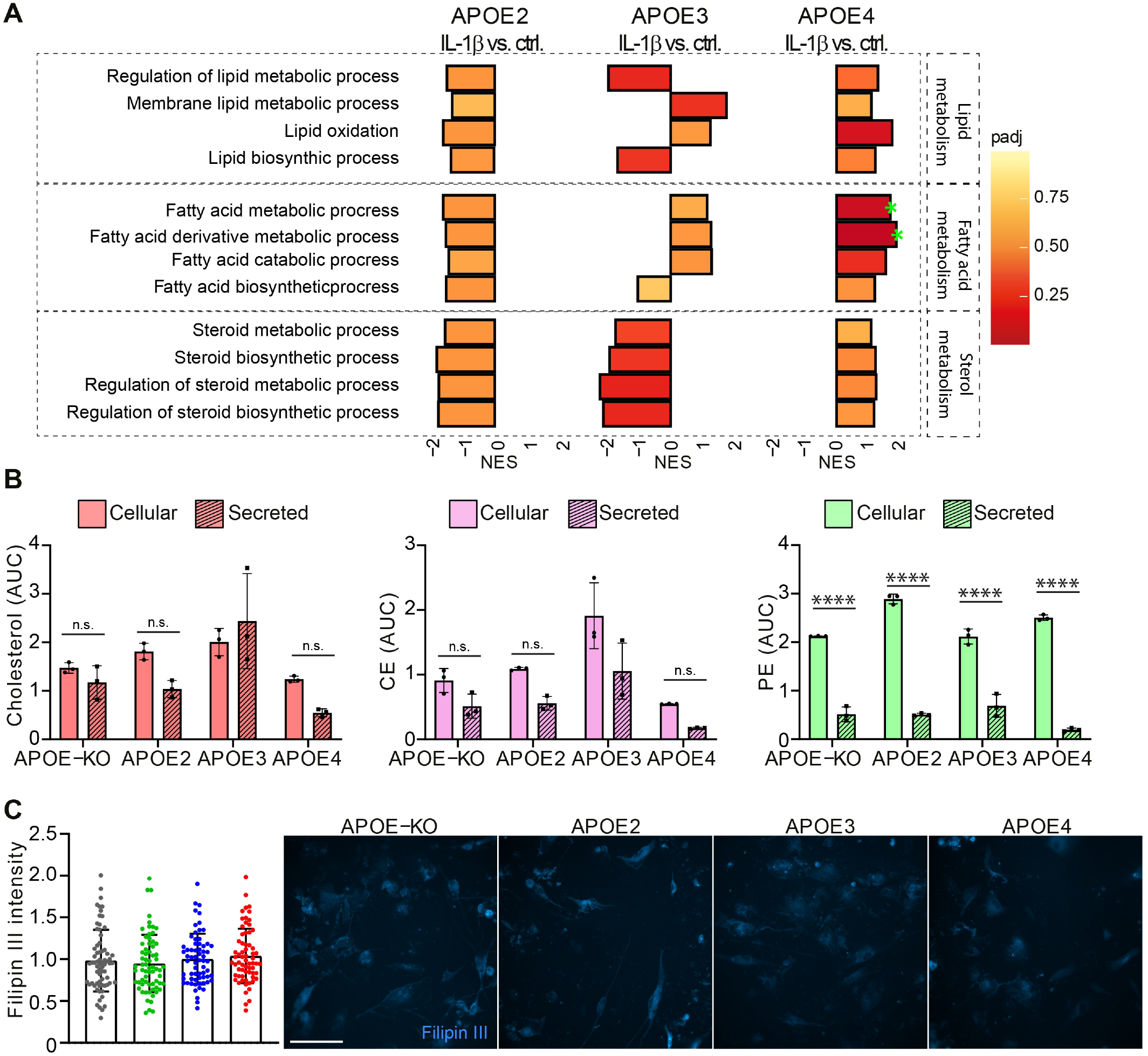
IL-1β treatment induces lipid, fatty acid and sterol metabolism in APOE4 iAstrocytes. A. GSEA of differential protein expression at baseline vs IL-1β-treated iAstrocytes (APOE2, E3 and E4 respectively). Normalized enrichment score (NES) is plotted on the x-axis, with color-coaded bars according to adjusted p-value (Padj). Padj 0.2 was considered significant, annotated with green stars. B. Cellular and secreted lipids determined with HPTLC, normalized to cellular phosphatidylcholine. C. Bar graphs of Filipin Ill intensity quantification, normalized to APOE3 intensity, with representative example images on the right side. Scale bar: 150μm. Data represent mean± SD.(****: p < 0.0001, Two-way ANOVA with post hoc Holm-Šídák’s multiple compari sons test)

## Discussion

Numerous methods deriving astrocytes from iPSCs have been successfully developed over the past years (de Leeuw and Tackenberg, 2019). With establishing an iPSC-astrocyte model to study AD-related APOE biology, it was essential that the astrocytes recapitulated a mature resting state, as they do in the human brain (Zhang et al., 2016). Even though we do use FCS in the initial stage of astrocyte differentiation, we inhibit proliferation with cytosine arabinoside and withdraw FCS for the last two weeks before analysis, slowing down cell division. Research focussing on the transcriptomic profile of activated astrocytes *in vitro*, is mostly based on rat or mouse studies (Zamanian et al., 2012), and it is yet unclear whether transient exposure of human astrocytes to FCS will impose a persistent activated state. We show evidence that our iAstrocytes drastically change in morphology from day 30 (with FCS), to day 44 (without FCS), in combination with the drastically reduced proliferation rate, providing evidence that even with previous FCS exposure, our iAstrocytes further progress into a mature resting state. In addition, iAstrocytes can be stimulated with low levels of pro-inflammatory cytokine, relevant in AD, whereas day 30 astrocytes derived with this protocol were previously activated with higher concentrations of stimuli (TCW *et al*., 2017). Additionally, the same study has shown activation of astrocytes by treatment with Aβ42 peptide. Interestingly, our day 44 iAstrocytes are not susceptible to activation by Aβ42, possibly pointing out that these cells need to be primed in order to be reactive to Aβ. These results may provide novel insights to the order of the cellular phase of AD progression, where we suggest an inflammatory microenvironment preceding or being necessary for Aβ42 stimulation of the cells (De Strooper and Karran, 2016).

### Homeostatic functions

Astrocytes play a pivotal homeostatic role in the brain, providing neural cells with cell membrane components such as lipids and cholesterol, taking up excess of neurotransmitters at the tripartite synapse to prevent excitotoxicity, and phagocytosing cellular debris and waste, alongside microglia (Zorec et al., 2017). Many of these functions have been found to be altered or dysfunctional in AD, such as a decrease in glutamate uptake (Scott et al., 2011). We show that *APOE* has an allele-dependent effect on glutamate uptake. *APOE-KO* and *APOE2* showed similar uptake capacity, whereas *APOE3* and *APOE4* iAstrocytes displayed lowest uptake. Interestingly, *APOE4* is associated with an increased risk for and an earlier onset of chronic temporal lobe epilepsy (Briellmann et al., 2000; Li et al., 2016). Further, AD patients are at higher risk for developing seizures and epilepsy (Pandis and Scarmeas, 2012). As elevated glutamate levels are linked to seizures (Barker-Haliski and White, 2015), the reduced glutamate uptake capacity in *APOE4* iAstrocytes may be a major contributing factor to the pathology and onset of seizures in AD as well as temporal lobe epilepsy. Previous research on humanized APOE mice showed reduced glutamine levels, and reduced capacity to incorporate glutamate into the brain of *APOE* 4TR mice, while glutamate transporters were not significantly changed (Dumanis et al., 2013). APOE4 also interacts with insulin receptors and other transporters, relocating them to the endosomal compartment: a similar mechanism may also affect glutamate transporters EAAT and GLAST surface expression, ultimately decreasing glutamate uptake capacity in the *APOE4* iAstrocytes (Yassine and Finch, 2020).

A similar pattern was detected in Aβ42 uptake (KO>E2>E3=E4), which is likely facilitated by APOE-bound Aβ42 clearance; a complex formation that is genotype dependent (E2>E3>E4) (Aleshkov et al., 1997). Interestingly, *APOE-KO* iAstrocytes were most proficient in taking up Aβ42, probably through direct lipoprotein receptor uptake of Aβ42 (Basak et al., 2012). *APOE*-dependent Aβ42 uptake may be the primary mode of degradation, even though an *APOE*-independent route, as seen in *APOE-KO* iAstrocytes, might be more efficient. Aβ42 uptake was reduced by an LRP1 antagonist in all genotypes except *APOE4*. However, LRP1 did not show a genotype-dependent difference in total protein level. In line with these results, LRP1 surface expression (but not total protein), was found to be decreased in *APOE*4 mouse astrocytes, caused by dysfunctional endosomal recycling of LRP1 to the cell surface (Prasad and Rao, 2018). This mechanism could explain our findings of reduced LRP1-mediated Aβ42 uptake in *APOE4* iAstrocytes. Even though we did not observe any endosomal deficit by assessing RAB5 expression, endosomal pH and other functional measures could still affect receptor recycling.

Altogether we provide evidence that the presence of the *APOE*4 allele induces a gain-of(toxic)-effect. *APOE2* and *APOE-KO* iAstrocytes show a similar phenotype, indicating differential homeostatic mechanisms by different APOE isoforms or lack thereof, in human iAstrocytes. The *APOE* allele-dependent effect on glutamate uptake and LRP1-dependent Aβ42 uptake can provide a mechanism explaining the biological role of *APOE* in homeostatic functions such as regulation of glutamate levels, but also pathological events such as extracellular Aβ42 build up.

### Lipid and cholesterol metabolism and the endolysosomal system

GSEA of the proteomic data set revealed downregulation of lipid/sterol metabolism and biosynthesis in *APOE4* compared to *APOE3* and *APOE2* iAstrocytes, whereas lipid catabolic processes show opposite effects. Triacylglycerols are subject to lipid catabolism, where they are broken down into glycerol and fatty acids, followed by fatty acid beta oxidation (Edwards and Mohiuddin, 2021) - another pathway upregulated in *APOE4* iAstrocytes. Our HPTLC results showed an allele-dependent increase in TAGs, indicating that lipid accumulation may lead to an imbalance in lipid metabolism/biosynthesis and catabolism, which are counter regulated in *APOE2* and *APOE4* iAstrocytes. These results are in line with a recent study, showing increased lipid content and decreased fatty acid oxidation in *APOE*4 mouse astrocytes (Qi et al., 2021).

Cholesterol and cholesteryl esters (CEs), however, showed an allele dependent decrease in cellular content (E2=E3>E4=KO), and efflux (E2=E3=KO>E4), in line with our proteomic findings. The first committed enzyme in *de novo* biosynthesis of cholesterol, squalene synthase (FDFT1), showed very strong allele-dependent downregulation (E4>E3>E2=KO), indicating that indeed these cells have a decreased capacity to produce cholesterol (Do et al., 2009). At the same time ABCA1 also showed allele-dependent expression levels (KO=E2>E3>E4). This protein is vital in facilitating cholesterol efflux and is upregulated in a positive feedback loop via liver X receptor (LXR), or LXR-independent cholesterol efflux, as well as APOE lipidation and production of cholesterol-rich HDL particles (Fan et al., 2018; Oram, 2003). The decrease of ABCA1 could explain why there is no significant decrease in cellular cholesterol and CEs in *APOE-KO* and *APOE4* Astrocytes, but there is a significant decrease of efflux in the latter genotype. This again indicates, that in the presence of APOE, there is a primary APOE-dependent mechanism of efflux, but in its absence, cholesterol or CEs may be secreted through passive efflux, APOA1-dependent ABCA1 efflux, or ABCGR1-dependent relocation of cholesterol to the surface, becoming available to HDL particles (Sharma and Agnihotri, 2019). Additionally, the excess TAG binding and occupying APOE4 can leave the lipoprotein unable to aid in cholesterol efflux. Even though an APOE-independent mode of efflux might be possible, *APOE2* and *APOE3* iAstrocytes show significantly higher levels of cellular cholesterol and CEs, indicating that E2 or E3 are necessary to maintain adequate levels of cellular cholesterol and CEs. Another study showed that APOE4 and ABCA1 co-aggregate, reducing ABCA1 recycling to the membrane (Rawat et al., 2019). We further show that also total levels of ABCA1 are decreased in *APOE4* iAstrocytes, suggesting it is not solely a mechanistic dysfunction, but decreased protein level altogether.

The increase in non-membrane bound cholesterol in *APOE3* and *APOE4* iAstrocytes (E4>E3>E2=KO) is in line with the decrease in cholesterol efflux; membrane bound cholesterol can bind to HDL-particles and be secreted, and APOE-mediated efflux may be ineffective in *APOE4* due to the lack of ABCA1. APOE4-bound cholesterol, taken up by the cell, is released from the receptor in the late endosome or lysosome and aggregation of APOE4 has been shown to cause endosomal congestion and dysfunction (Morrow et al., 2002; Narayan et al., 2020; Nuriel et al., 2017; Rawat *et al*., 2019). We indeed observed an increased colocalization of non-membrane bound cholesterol to lysosomes (E4>E3=KO>E2).

Although we did not detect significant changes in RAB5 protein levels or cell area covered, it does not exclude the presence of functional changes or it may require prior activation of the astrocytes to detect endosomal deficits, which would thus not be present in our cells at baseline. However, we did observe a decrease in LAMP2 protein levels (E2=E4=KO>E3). Remarkably, *APOE3* iAstrocytes displayed the highest levels, and other genotypes showed similar lower levels. In contrast, the percentage of cell area covered by LAMP2 was not significantly changed. It is known that a decrease in LAMP2 levels cause a lack of lysosomal trafficking and decrease in autophagic flux (Cui et al., 2020). However, the decrease of LAMP2 in itself may not be suggestive of a pathological mechanism, as *APOE-KO* and E2 iAstrocytes displayed similar LAMP2 protein levels to E4 cells. These results indicate that the presence of APOE3 and more so APOE4 play a role in rerouting cholesterol to the lysosomes, although *APOE4* lysosomes may not be equipped to process excess material due to low lysosomal flux. Additionally, *APOE2* astrocytes showed least colocalization of non-membrane bound cholesterol to lysosomes, even less than *APOE-KO* iAstrocytes. This indicates that APOE2-mediated cholesterol flux is more efficient than compensatory mechanisms taking over in the absence of APOE.

### Cholesterol metabolism and inflammatory regulation

Immune regulatory pathways in iAstrocytes at baseline were negatively enriched in *APOE*2 vs E3, highly positively enriched in *APOE*4 vs E3 and moderately enriched in *APOE*4 vs E2. Two pathways related to TGF-β signalling were opposingly regulated (down in *APOE*4), in line with a study showing dysfunction in TGF-β signalling in models of early AD and in human AD brain (Tesseur et al., 2006), likely connected to its neuroprotective role. Additionally, TGF-β can be directly regulated by APOE in microglia with a neurodegenerative phenotype (Krasemann et al., 2017). Generally, mild immune regulation in *APOE*4 cells ensures the capacity for an increase in response after activation, which we observed after IL-1β stimulation: *APOE*2 showed negative enrichment, *APOE*3 positive enrichment while *APOE*4 iAstrocytes displayed significant negatively or positively enriched pathways. Additionally, more pathways were affected in total (both up and down) in *APOE*4 iAstrocytes than in *APOE*3 and *APOE*2, indicating that *APOE*4 iAstrocytes are more susceptible to inflammatory stimuli. Interestingly, two of the pathways downregulated in *APOE*4 iAstrocytes were NFkB and IL-1 mediated signalling, which were not significantly regulated in *APOE*3 or E2. Many studies show an increase in NFkB in AD brain or mouse models of AD, however, our human iAstrocytes are at a resting state prior to IL-1β treatment, and thus do not model reactive astrocytes or late-stage AD (Rangaraju et al., 2018). On the contrary, neuroregenerative mechanisms in Zebrafish models of AD have shown a decrease in NFkB signalling (Bhattarai et al., 2020). It is possible that neuroprotective mechanisms compensate for the general inflammatory phenotype of the iAstrocytes, which is evident by downregulated NFkB and IL-1 mediated signalling pathways in *APOE*4 iAstrocytes. On a functional level, we observed a strong allele-dependent increase in inflammatory cytokine release upon iAstrocyte activation (E4>E3>E2>KO), and even IL-6 and IL-8 release from *APOE*4 at baseline. This indicates that *APOE*4 iAstrocytes show some reactive capacity already at baseline, which is exacerbated by IL-1β treatment. This is in line with research showing that chronic inflammation is associated with an earlier onset of AD in *APOE*4 carriers. Our findings of an inflammatory phenotype in resting *APOE*4 iAstrocytes could explain the mechanistic link between chronic inflammation and AD pathogenesis in *APOE*4 carriers (Tao et al., 2018).

Activation of iAstrocytes also resulted in an increase in pathways classified as lipid, fatty acid, and cholesterol metabolic processes in *APOE*4 iAstrocytes while the opposite effect was seen in *APOE*2 iAstrocytes. *APOE*3 cells showed a decrease in sterol metabolic pathways upon activation while lipid and fatty acid metabolism were not regulated uniformly. These results oppose the findings at baseline. The significant difference between intracellular and secreted cholesterol in *APOE*4 iAstrocytes was abolished after IL-1β treatment, but had no effect on CE or PE, indicating that IL-1β treatment specifically affects cholesterol homeostasis.

It has been shown that cytokine stimulation can regulate cholesterol synthesis by activating SREBP1 (Gierens et al., 2000). This transcription factor in turn induces *de novo* biosynthesis of cholesterol and increases the expression of LRP1 or VLDLR to increase cholesterol uptake (Shimano, Nature rev end, 2017). However, cellular cholesterol did not show an increase upon IL-1β treatment in *APOE*4 iAstrocytes. Cholesterol biosynthesis and lipid efflux machinery are regulated via LXR and RXR complex activation; when intracellular cholesterol is increased, LXR/RXR increases ABCA1 expression to induce lipid efflux, APOE expression and inhibition of SREBP activation (Oram, 2003; Shimano and Sato, 2017). Our results show an allele-dependent regulation of cellular cholesterol (E4=KO>E3<E2), with more cholesterol being translocated into the lysosomes in *APOE*4 iAstrocytes, which could lead to a lower degree of LXR activation. LXRs also inhibit NFκB signaling (Bi and Song, 2016), but this pathway is shown to be reduced in *APOE*4 iAstrocytes, indicative of an inherent dysfunction in the signalling complex. The LXR/RXR complex is in turn linked to transcription factor PPARy, which can also induce ABCA1 and APOE expression. In line with our results, PPARy signalling was decreased in *APOE*4 transgenic mouse brain, and increased in *APOE*2 (Wu, 2018). Furthermore, genetic variability of LXR may contribute to the risk for AD, in addition to the ability of LXR agonists to attenuate Aβ pathology (Kang 2012). Hence there may be deficits in the metabolically linked PPARy and LXR/RXR complex, rendering them less potent to induce cholesterol upregulation upon IL-1β activation in *APOE*4 iAstrocytes, as seen in our results. However, LXR/RXR are likely not entirely dysfunctional as cellular/secreted cholesterol is not significantly different after IL-1β treatment in contrast to baseline, suggesting residual LXR activity allowing for the relative increase in efflux of cholesterol.

In addition, IL-1β treatment abolished the allele-dependent non-membrane bound increase in cholesterol load that was observed at baseline by Fillipin III staining. Considering the significant increase in cellular *APOE*3 cholesterol (and a non-significant increase in *APOE*2) upon IL-1β treatment, which was not present *APOE*4, we conclude that the effect of IL-1β seen is due to an increase of total cholesterol in the *APOE*2 and E3 iAstrocytes, but not in *APOE*4.

Cholesterol metabolism and immune regulation were also shown to be linked through ABCA1, ABCGR1 and HDL particles (Yvan-Charvet et al., 2010). ABCA1- or ABCGR1-deficiency increased inflammatory activation through excess of cholesterol in the membranes, which anchored Toll-Like Receptors. Although we found an allele-dependent decrease in ABCA1, we did not observe an allele-dependent increase in total cholesterol, rather the opposite: *APOE*4 iAstrocytes have the least amount of total cellular cholesterol, but more non-membrane bound cholesterol (Fillipin III) than E2, E3 and KO cells. This indicates that the link between cholesterol metabolism and immune regulation is likely regulated by nuclear receptors as mentioned previously.

Together, we show that *APOE*2, E3, E4 and KO iAstrocytes show distinct phenotypes in homeostatic functions, cholesterol and lipid metabolism, lysosomal function and inflammatory regulation. While several cellular functions, such as glutamate uptake, seem to display a clear allele-dependent decrease in capacity, the interplay between cholesterol metabolism and immune regulation indicates that it is not a mere decrease of function in *APOE*4 cells: APOE2, APOE3 and APOE4 iAstrocytes may utilize different mechanisms altogether. Further mechanistic insights into the difference in APOE2 and E4 iAstrocytes are imperative to continue the understanding of fundamental APOE biology, which simultaneously proves its inherent relevance in Alzheimer’s disease. Our human isogenic mature iAstrocyte cell model represents a powerful tool to further address these open questions.

## Materials and Methods

### Cells

iPSC lines were purchased from EBISC (BIONi010-C-2, C-3, C-4 and C-6), harbouring a single *APOE*2, *APOE*3 or *APOE*4 allele generated using CRISPR, or an *APOE* knockout generated by an insertion-deletion mutation in exon 2 (Schmid *et al*., 2019; 2020).

### iPSC differentiation to neural progenitor cells

iPSCs were maintained on Vitronectin (StemCell Technologies) coated plates in TesRE8 medium (StemCell Technologies) and passaged using ReLeSR (StemCell Technologies). iPSCs and subsequent cell types were frozen in BAMBANKER™ serum-free cryopreservation medium (Wako Chemicals). Neural progenitor cells were derived as described (Sancho-Martinez et al., 2016) with minor adjustments. iPSCs were dissociated with Accutase (Sigma-Aldrich), and passaged onto Vitronectin-coated plates, at 80’000 cells/well of a 12-well plate, in TesRE8 medium + 2µM Thiazovivin. Within 24hrs the cells were washed with PBS (Sigma-Aldrich), and changed with TesRE8 medium. On day 0 cells should be approximately 20% confluent, were washed with PBS and changed with Neural Induction Medium 1 (NIM-1: 50% Advanced DMEM/F12 (Invitrogen), 50% Neurobasal (Invitrogen), 1x N2 (Gibco), 1x B27 (Gibco), 2 mM GlutaMAX (Gibco) and 10ng/mL hLIF (Peprotech), 4mM CHIR99021 (Miltenyi), 3mM SB431542 (Miltenyi), 2mM Dorsomorphin (Miltenyi) and 0.1mM Compound E (StemCell Technologies). Medium was changed daily for 2 days, then cells were washed with PBS and switched to Neural Induction Medium 2 (NIM-2: NIM-2: 50% Advanced DMEM/F12, 50% Neurobasal, 1x N2, 1x B27, 2mM GlutaMAX, 10ng/mL hLIF, 4mM CHIR99021, 3mM SB431542 and 0.1mM Compound E). After another 4 days, cells were approximately 80% confluent and passaged at 1:4 from 12-well plates to 15µg/mL Poly-L-Ornithine (Sigma-Aldrich) + 10µg/mL Laminin (Sigma-Aldrich) coated 6-well plates, in Neural Stem cell Maintenance Medium (NSMM: 50% Advanced DMEM/F12, 50% Neurobasal, 1x N2, 1x B27, 2mM GlutaMAX, 10ng/mL hLIF, 3mM CHIR99021 and 2mM SB431542). Medium was changed every day, and NPCs were kept at high density: passaged once 90-100% confluent, approximately 2x a week. 2µM Thiazovivin was added to the medium for the first 7 passages, and after 5 passages 5ng/mL FGF (ThermoFisher) + 5ng/mL EGF (ThermoFisher) was added to NSMM. For assays, NPCs were plated at 200’000 cells per well of a 24-well plate.

### NPC differentiation to iAstrocytes

Between passage 1-4, NPCs were differentiated to astrocytes by the application of Primary Astrocyte Medium (AM; ScienCell: 1801, astrocyte medium (1801-b), 2% FCS (0010), for 30 days according to TCW and colleagues (TCW *et al*., 2017). NPCs were plated on 1µg/mL Fibronectin (Sigma-Aldrich) coated plates, at 135’000 cells/well of a 6-well plate in NSMM. After 24hrs the cells were washed with PBS and changed to AM. Medium was changed every other day, and cells were split at initial density once 80-100% confluent. To obtain mature, non-proliferative cells, astrocytes were replated on day 30 to 350’000 cells/well of a 6-well plate, in AM without FCS + 2µM AraC (AM+). Medium was changed every other day; cells were not actively proliferating anymore so no passaging was needed. At day 37 the astrocytes were washed with PBS and medium replaced with AM without FCS, and without AraC (AM-). Astrocytes were kept in AM-until day 44 and subsequently plated for experiments accordingly: µ-Slide Angiogenesis slides (Ibidi), 10’000 cells/well; 96-well plates, 15’000 cells/well; 24-well plates + coverslips, 80’000 cells/well; 24-well plates: 200’000 cells/well, 6-well plates: 350’000 cells/well. Cells were always plated 24 hours prior to treatment or assay.

### Immunocytochemistry

Astrocytes were grown on coverslips for 1-2 days and fixed with 4% Paraformaldehyde (Electron Microscopy Science) in PBS + 4% Sucrose (Sigma-Aldrich) for 20min at room temperature (RT), and washed 3x with PBS. Cells were blocked in 10% donkey serum with 0.1% Triton X-100 in PBS for 1hr at RT, and subsequently incubated with primary antibody in 3% donkey serum with 0.1% Triton X-100 in PBS overnight at 4°C. Primary antibodies were used accordingly; S100β (Sigma, S2532, 1:200), GJA1 (Abcam, ab235585, 1:200), GFAP (Sigma, G9269, 1:200), Rab5 (Abcam, ab218624, 1:1000), Lamp2 (Biolegend, 354302, 1:100), amyloid-β 1-42 (Invitrogen, 44-344, 1:1000). Cells were washed 3x with PBS and incubated with the respective secondary antibody at 1:500 in 3% donkey serum with 0.1 % Triton X-100 in PBS for 2hrs at RT (anti-rabbit Alexa Fluor 568 and anti-mouse Alexa 488 both from donkey, Jackson Immunoresearch). Cells were washed 1x and incubated with 0.2µg/mL DAPI (4’,6-diamidino-2-phenylindole, Sigma-Aldrich) for 10min at RT. Cells were washed 2x with PBS before mounting with Mowiol® 4-88 (Sigma-Aldrich).

### Protein sample preparation

Cells were dissociated at day 44-46 and lysed in RIPA buffer (50 mM Tris-HCL pH7.6, 150 mM NaCl, 1% NP40, 0.5% sodium dodecyl sulfate, 0.5% sodium deoxycholate, and 2mM EDTA (Sigma)) supplemented with cOmplete^™^ Protease Inhibitor Cocktail (Roche). Lysates were incubated on ice for 20min, followed by sonication and centrifugation (Eppendorf), at 10’000 x g for 10min at 4°C. Total protein concentration was determined using a Pierce BCA Protein Assay Kit (ThermoScientific).

### Western blot

Protein samples were loaded onto a Novex™ 10 to 20%, Tricine, 1.0 mm, Mini Protein Gel (Thermo fisher), and transferred semi-dry onto a Mini 0.2µm Nitrocellulose blot (Biorad). Blots were blocked with 5% non-fat milk (AppliChem) in PBS, and subsequently stained with primary antibody overnight in 5% non-fat Milk in PBS + 0.05% Tween 20 (Sigma). The following primary antibodies were used; FDFT1 (Abcam, ab195046, 1:2000), ABCA1 (Abcam, ab18180, 1:500), HMG-CoA Reductase (Merck-Millipore, ABS229, 1:700), LRP1 (Abcam, EPR3724, 1:1500), LXR (Abcam, ab24362, 1:500), S100α6 (Abcam, ab181975, 1:1000), α-tubulin (Sigma, T9026, 1:1000), β-actin (Abcam, ab6276, 1:20’000), GJA1 (Abcam, ab235585, 1:1000), Rab5 (Abcam, ab218624, 1:1000), Lamp2 (Biolegend, 354302, 1:500). Blots were subsequently incubated with peroxidase-labeled secondary antibody (anti-goat; 705-036-147, anti-mouse; 715-035-151, anti-rabbit; 111-035-144, Jackson ImmunoResearch, 1:5000), and developed with ECL Prime or Select (Amersham), or Pierce™ ECL (ThermoScientific).

### Glutamate uptake assay

Astrocytes were seeded onto a 20μM fibronectin-coated 96-well plate at previously described density. After 24hr, medium was changed with DMEM without glutamine (Gibco), for 6hr. Subsequently, cells were washed with HEPES buffer (25mM HEPES, 140mM NaCl, 3.5mM KCl, 2.5mM CaCl_2_, 1mM MgCl_2_, 0.1% BSA, pH 7.5; Sigma), and incubated for 60min with 20μM glutamate (Sigma) at 37°C. After incubation the residual glutamate was transferred to a white Nunc™ F96 MicroWell™ Polystyrene Plate (ThermoScientific), and kept on ice. The cells were washed with PBS and directly lysed in RIPA buffer, following 20min incubation on ice. During this time the Glutamate Detection Reaction from the Glutamate-Glo assay (Promega) was added to the residual glutamate, and incubated for 60min at room temperature according to protocol. At this time the cell lysates were centrifuged for 15min at 4000 x g at 4°C, following total protein measurement using the Pierce BCA Protein Assay Kit (ThermoScientific). Residual glutamate was measured with luminescence using a TECAN Infinite M1000 plate reader. Glutamate uptake was measured by subtracting the residual glutamate from the positive control, and normalized by the total protein content.

### Meso Scale Discovery cytokine measurement

Astrocytes or NPCs were plated on 24-wells plates, 24hrs prior to treatment. Both cell types were incubated with medium only, additionally astrocytes were treated with 10ng/mL IL-1β or TNF-α (StemCell Technologies), for 24hr. Supernatant was transfered to fresh Eppendorf Tubes, and cells were harvested for protein samples and used in label-free mass spectrometry. Supernatant was centrifuged at 21’000 x g for 5min at 4°C, and diluted at 1:4 in Diluent 2 from the MSD^®^ MULTI-Spot Assay system, pro-inflammatory panel 1 (human) kit. With this kit 10 cytokines were measured; IFN-γ, IL-1β, IL-2, IL-4, IL-6, IL-8, IL-10, IL-12p70, IL-13, and TNF-α, according to protocol.

### Mass spectrometry proteomic analysis

Astrocytes were replated to 24-well plates, 24hrs prior to treatment. Cells were treated as mentioned in the previous paragraph, and cells were lysed in RIPA buffer as described. Samples were further processed by using a commercial iST Kit (PreOmics, Germany) with an updated version of the protocol. Briefly, 100 μg of proteins were solubilized in ‘Lyse’ buffer, boiled at 95°C for 10min and processed with High Intensity Focused Ultrasound (HIFU) for 30s setting the ultrasonic amplitude to 85%. The samples were digested by adding 10μl of the ‘Digest’ solution. After 60 min of incubation at 37°C the digestion was stopped with 100μl of Stop solution. The solution was transferred to the cartridge and were removed by centrifugation at 3800 x g, while the peptides were retained by the iST-filter. Finally, the peptides were washed, eluted, dried and re-solubilized in 20μl of 3% acetonitrile, 0.1% FA. iRT peptides (Biognosys) were added to each vial.

Mass spectrometry analysis was performed on an Orbitrap Fusion Lumos (Thermo Scientific) equipped with a Digital PicoView source (New Objective) and coupled to a M-Class UPLC (Waters). Solvent composition at the two channels was 0.1% formic acid for channel A and 0.1% formic acid, 99.9% acetonitrile for channel B. For each sample 2μl of peptides were loaded on a commercial MZ Symmetry C18 Trap Column (100Å, 5µm, 180µm x 20 mm, Waters) followed by nanoEase MZ C18 HSS T3 Column (100Å, 1.8µm, 75µm x 250mm, Waters). The peptides were eluted at a flow rate of 300nl/min by a gradient from 5 to 22% B in 80 min and 32% B in 10 min after an initial hold at 5% B for 3 min. The column was washed with 95% B for 10min and afterwards the column was re-equilibrated to starting conditions for additional 10 min. Samples were acquired in a randomized order. The mass spectrometer was operated in data-dependent mode (DDA) acquiring a full-scan MS spectra (300− 1’500 m/z) at a resolution of 120’000 at 200 m/z after accumulation to a target value of 500’000. Data-dependent MS/MS were recorded in the linear ion trap using quadrupole isolation with a window of 0.8 Da and HCD fragmentation with 35% fragmentation energy. The ion trap was operated in rapid scan mode with a target value of 10’000 and a maximum injection time of 50ms. Only precursors with intensity above 5’000 were selected for MS/MS and the maximum cycle time was set to 3 s. Charge state screening was enabled. Singly, unassigned, and charge states higher than seven were rejected. Precursor masses previously selected for MS/MS measurement were excluded from further selection for 20s, and the exclusion window was set at 10ppm. The samples were acquired using internal lock mass calibration on m/z 371.1012 and 445.1200. The mass spectrometry proteomics data were handled using the local laboratory information management system (Türker et al., 2010).

### Proteomics data processing and analysis

The acquired raw MS data were processed by MaxQuant (version 1.6.2.3). We obtained protein identification using the integrated Andromeda search engine. We used the canonical sequence Uniprot FASTA database (organism ID 9606, proteome ID UP000005640) downloaded from uniprot.org in July 2019, concatenated to its reversed decoyed database and common protein contaminants (http://fgcz-proteomics.uzh.ch/fasta/fgcz_9606_reviewed_cnl_20190709.fasta). Carbamidomethylation of cysteine was set as fixed, while methionine oxidation and N-terminal protein acetylation we specified as variable modifications. Enzyme specificity was trypsin/P, allowing a minimal peptide length of 7 amino acids and a maximum of two missed cleavages. For search and label-free quantification (LFQ), we used MaxQuant Orbitrap default settings. The maximum false discovery rate (FDR) was set to 0.01 for peptides and 0.05 for proteins. For LFQ, we specified a 2-minute window for match-between-runs. In the MaxQuant experimental design template, each file is kept separate in the experimental design to obtain individual quantitative values.

We inferred protein intensity estimates from peptide intensity values reported in the MaxQuant generated peptides.txt file. We preprocessed the peptide intensities by removing intensities equal to zero and log2 transforming non-zero intensities. To remove systematic differences between samples, we applied a modified robust z-score transformation that preserves the data’s original variability. We obtained protein intensity estimates by fitting Tukey’s median polish to the peptide data. To estimate fold changes among conditions, i.e., cell lines or APOE genotypes, we fitted a linear model to each protein, then calculated contrasts, and used the Wald test to obtain p-values. Leveraging the parallel structure of the high throughput experiment, we used experimental Bayes to moderate protein variance estimates and updated the t-statistics and p-values accordingly [Smyth, Gordon K]. The p-values were adjusted for multiple testing, using the Benjamini Hochberg procedure and we obtained false discovery rates (FDR). We used the implementation of these methods available in the R package prolfqua (Wolski et al., 2020).

We performed gene set enrichment analysis using the R/Bioconductor package fgsea (Korotkevich et al., 2021) and used gene sets specified in the molecular signature database (http://www.gsea-msigdb.org/). To apply GSEA to proteomics data, we mapped the uniprot identifiers to Entrez Id’s using the UniProt mapping service. We ordered the protein lists using the moderated t-statistics. For cases where several UniProt Id’s were mapped to a single Entrez Id, we averaged the t-statistic (Smyth, 2004).

### Aβ uptake assay

Astrocytes were plated onto 24-wells plates, and treated with either 10ng/mL IL-1β, 0.2μM RAP (Molecular Innovations), or 10μM GSK2033 (Tocris), for 24hr. Lyophilized Hilyte488-tagged amyloid-β42 (Aβ42-Hilyte488, Anaspec) was diluted to 100μM and sonicated for 10min, followed by shaking at 300RPM for 24hrs at 4°C. After 24hr, cells were treated with the aforementioned compound, in combination with 1μM Aβ42-Hilyte488 for 2hr. Cells were then washed with PBS and and harvested with Trypsin-EDTA (Gibco), transferred to poly-propylene round bottom tubes (BD Bioscience) and washed with FACS buffer (20% FCS (Gibco), 5mM EDTA pH8, 0.01% NaN_3_ (Sigma), in HBSS (Gibco)). Cells were treated with 0.025% Trypan blue (Gibco) immediately before acquisition of the cells with the LSRII Fortessa 4L (BD Bioscience). Cells for example images were incubated with recombinant Aβ42 (rPeptide), and treated as mentioned before. Cells were incubated with 1μM Aβ42 peptide for 24hrs and fixed according to subchapter “immunocytochemistry”.

### Dextran-Alexa555- and Filipin III staining

Astrocytes were seeded onto µ-Slide Angiogenesis slides, and were additionally treated with Dextran-Alexa555 (ThermoFisher) if needed, otherwise cells were immediately used for Fillipin III staining. Incubation with 0.2mg/mL Dextran-Alexa555 was performed overnight, and followed by a 14hr chase period (changed to medium only to allow Dextran localization to the lysosomes). Astrocytes were then treated with 10mM Methyl-β-cyclodextrin for 30min, followed by fixation and treatment with the Cholesterol Assay kit (Abcam), according to protocol with minor adjustments. After fixation, astrocytes were incubated with Fillipin III for 2hr, followed by washing steps accordingly. Fillipin images alone were acquired with fluorescence microscopy, Dextra-Alexxa555 and Fillipin double labeling imaged with confocal microscopy.

### Immunofluorescence imaging and image processing

Confocal images were acquired with a Leica SP8 microscope, equipped with an HC PL APO CS2 20x objective (NA 0.75), using the Leica Application Suite X software. Fluorescent images were acquired with a Zeiss Axio Observer 7 equipped with a ORCA-flash4.9 camera and Plan-Apochromat 20x objective (NA 0.8), using the Zen 3.1 pro software. Image analysis and post processing was done in ImageJ or Imaris (Oxford instruments).

### Lipid extraction

Lipids from cell pellets and supernatants were extracted using a Bligh and Dyer protocol (Bligh and Dyer, 1959). Briefly, lipids were extracted by adding 3.8mL methanol:chloroform 2:1 (v:v) to a first Wheaton glass tube, followed by 1mL of sample: cell supernatant or cell pellet resuspended in HPLC-grade water. The tube was vortexed thoroughly. Then 1mL chloroform and 1mL water were added, followed by vortexing thoroughly after each addition and centrifugation at 2,000 rpm for 2min at 4°C. The lower phase was transferred to a second Wheaton tube, followed by the addition of 1mL chloroform and 1mL water. In parallel, 1mL chloroform was added to the first Wheaton tube and both tubes were vortexed thoroughly and centrifuged at 2000RPM for 2min at 4°C. The lower phase of the second Wheaton tube was transferred to a final Wheaton tube. The lower phase of the first Wheaton tube was transferred to the second Wheaton tube, vortexed thoroughly and centrifuged at 2000RPM for 2min at 4°C. The lower phase was added to the final Wheaton tube. The solvent was evaporated under vacuum at room temperature. Dried lipids were overlaid with argon and stored at -20°C until further analysis.

### High-performance thin-layer chromatography (HPTLC)

Lipids were spotted on HPTLC Silica Gel60 plates (Merck) using an automated TLC sampler (CAMAG ATS 4). The TLC plate was pre-washed in chloroform:methanol (1:1) and dried in a vacuum chamber. For neutral lipids separation and identification, a mix of lipid standards was used: 0.1μg cholesterol (Sigma), 0.1μg cholesteryl ester (Sigma, C9253), 1μg sphingomyelin (Sigma), 1μg phosphatidylcholine (Avanti, 850375), 1μg phosphatidylethanolamine (Avanti, 850725), 1μg triacylglycerol (Sigma). The lipid standard mix and conditioned media/cell pellets lipid extracts resuspended in chloroform:methanol:water (20:9:1) were spotted on the plate using the automated system. The plate was developed first in chloroform:methanol:ammonium hydroxide (65:25:4) for 5cm, dried briefly and re-developed in hexane:diethyl ether:acetic acid (80:20:2) for 9cm. After separation, the HPTLC plate was dried in a vacuum chamber for 30min. Lipids were visualized using a method adapted from Churchward and colleagues (Churchward et al., 2008). Briefly, a copper (II) sulfate staining solution was prepared: 5g of copper (II) sulfate (Sigma, 12849) dissolved in 40mL HPLC-grade water, filtered, mixed with 4.7mL of 85% ortho-phosphoric acid (Merck) and filled up to 50mL with HPLC-grade water. 10mL of freshly prepared staining solution was poured on the HPTLC plate, incubated for 1min, decanted and the plate was dried in the vacuum chamber for 15min. Lipids were then charred at 145°C for 7.5min. The plate was visualised in visible light, at 488nm and 546nm (ChemiDoc MP, Bio-Rad). Lipid spots were quantified using densitometry with the ImageJ software. Lipid content was normalized to cellular phosphatidylcholine content for each sample.

### Statistical analysis

Data are presented as mean ± standard deviation (SD). All datapoints (n-numbers) are plotted individually in each bar graph. Statistical analysis was performed with GraphPhad Prism (GraphPad Software Inc.) and RStudio. Proteomic datasets were analysed and visualized using edgeR (Robinson et al., 2010). All experiments were performed in triplicate, and a normality and lognormality test was performed, if necessary outliers were identified with the ROUT method (Q=1%). Significance was tested using Shapiro-Wilk test for normal distribution followed by Kruskal-Wallist test, with *post hoc* Dunn’s multiple comparisons test; or a one-way followed by *post hoc* Tukey’s multiple comparisons testing; or two-way ANOVA with *post hoc* Holm-Šídák’s multiple comparisons test. The statistical tests used in the different experiments are annotated in the respective figure legend.

## Acknowledgments

We thank Dr. Julia TCW, Mount Sinai Hospital, for advice and for providing reference cells to set up the astrocyte differentiation method. We are grateful to Stephanie Davaz and Debora Wanner, Institute for Regenerative Medicine, for cell culture support. We further thank the Functional Genomic Center and the Cytometry Facility of the University of Zurich for expert help on proteomic and flow cytometry analyses, respectively.

SMdL and CT were supported by the Betty and David Koetser Foundation for Brain Research and the Neuroscience Center Zurich. ACG and KL acknowledge the financial support of the Louis-Jeantet Foundation.

## Author Contributions

Conceptualization: SMdL and CT; Methodology: SMdL and CT; Software: RR and WW; Validation: SMdL, AWTK, KL; Formal analysis: SMdL, AWTK, KL, RR, WW and CT; Investigation: SMdL, AWTK, KL; Resources: ACG, RMN; Writing: SMdL and CT; Visualization: SMdL, RR, WW and CT; Supervision: ACG, RMN, CT; Project Administration: CT; Funding Acquisition: CT

---

**Figure S1.**
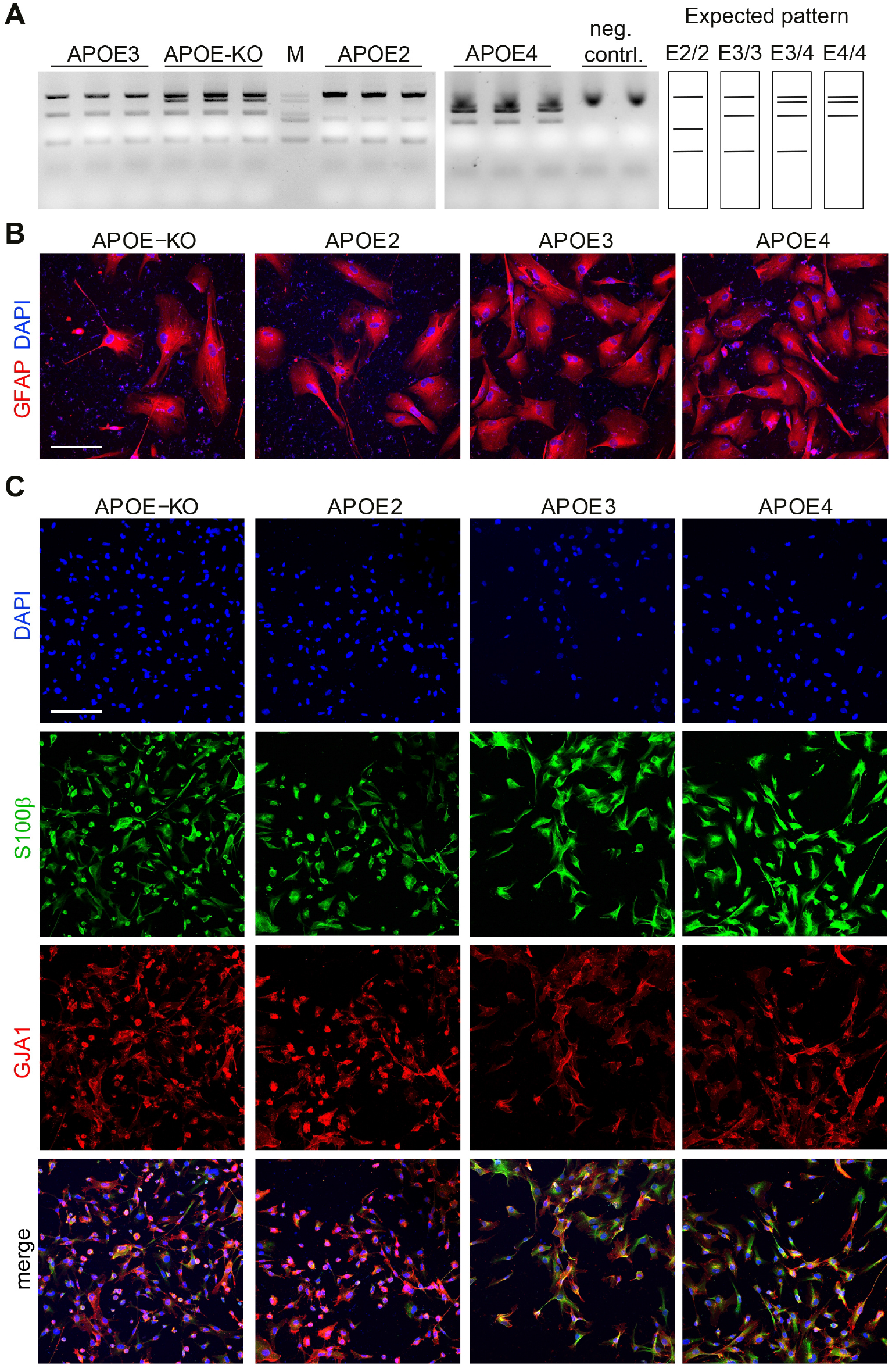
Genotyping and immunocytochemical characterization of APOE-isogenic iAstrocytes. A: Agarose gel of genotyping PCR using EzWay Direct ApoE Genotyping Kit with a schematic representation of the expected band pattern. B: Confocal images of iAstrocytes at day 44, stained with antibody against astrocyte marker GFAP. Scale bar: 150μm. C: Confocal images of iAstrocytes at day 30, stained with antibody against astrocyte markers S100β and GJA1. Scale bar: 150μm.

**Figure S2.**
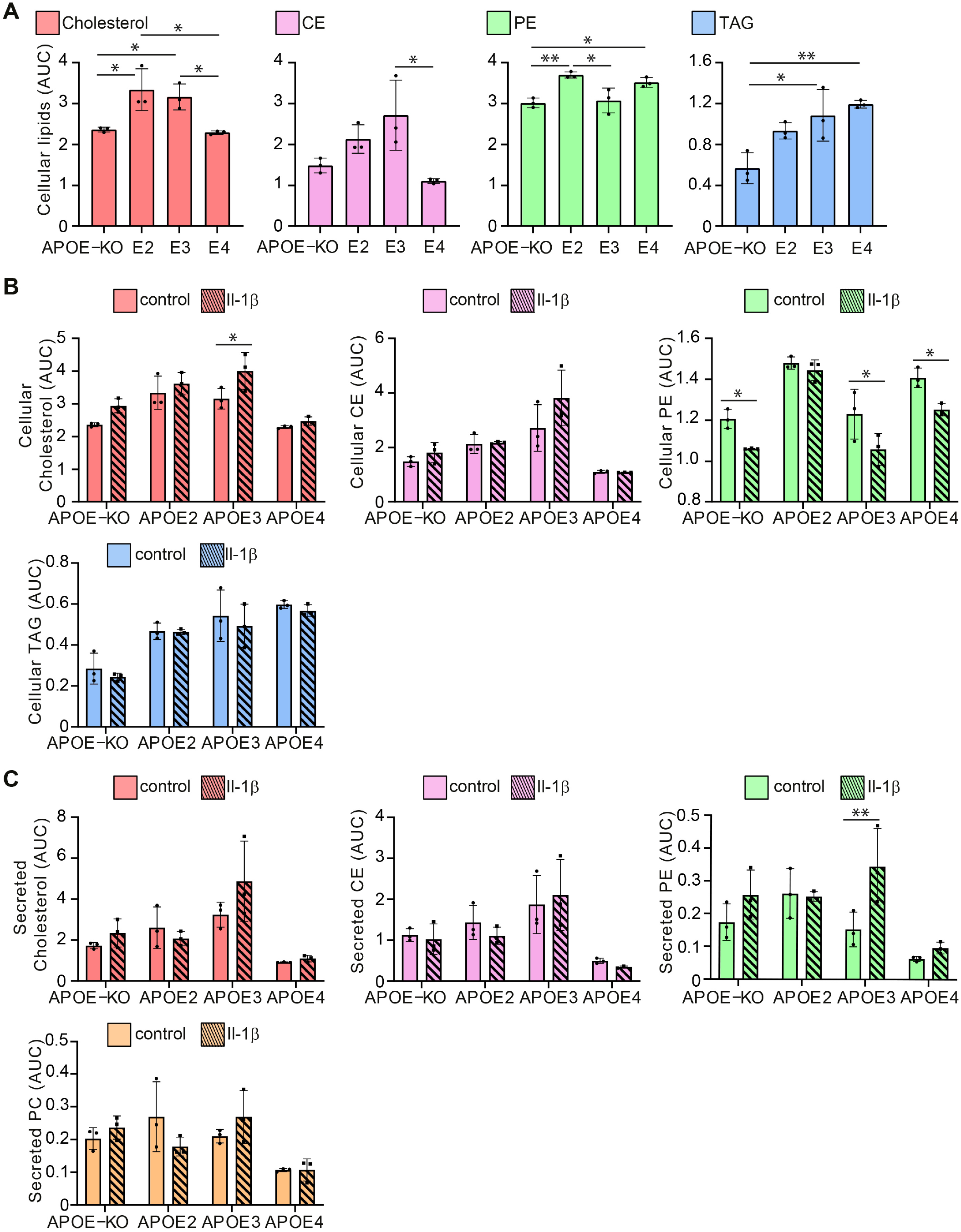
Cholesterol, CE and lipid analysis of the cellular and secreted fractions of iAstrocytes. A. Cellular cholesterol, cholesterylester (CE), phosphatydilethanolamine (PE), and triacylglycerol (TAG) quantified with HPTLC, normalized to cellular phosphatidylcholine (PC). B. Bar graphs of cellular cholesterol and lipid quantification at baseline and treated with 10ng/ml IL-1β. C. Bar graphs of secreted cholesterol and lipid quantification at baseline and after treatment with 10ng/ml IL-1β. Data represent mean± SD(*: p < 0.05; **: p < 0.01; ***: p < 0.001; ****: p < 0.0001, Two-way ANOVA with post hoc Holm-Šídák’s multiple comparisons test).

**Figure S3.**
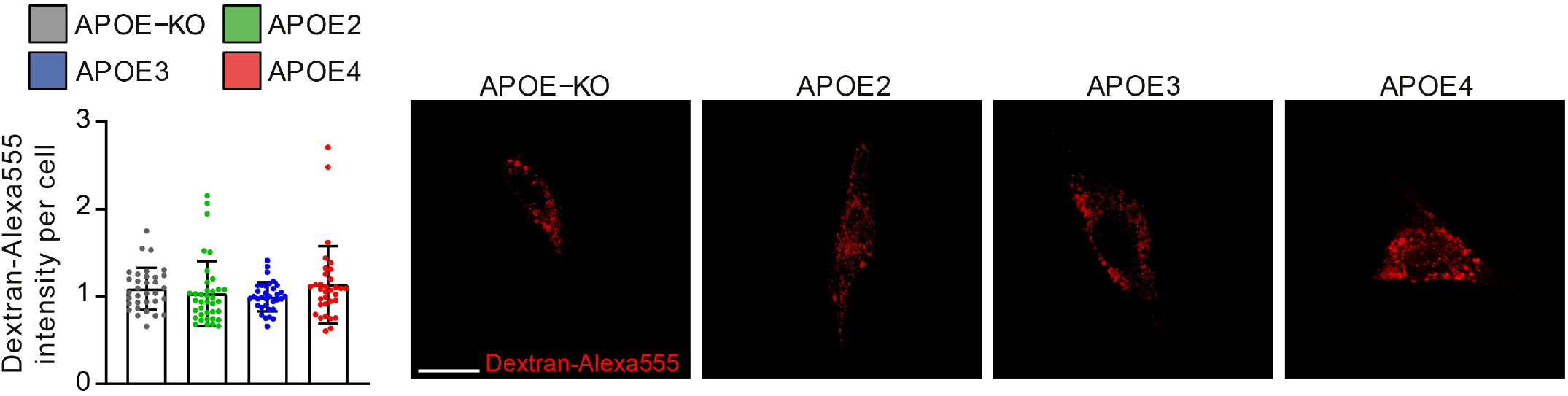
Dextran-Alexa555 uptake does not show genotype-dependent differences. Quantification of Dextran-Alexa555 intensity per cell, normalized to APOE3 iAstrocytes, (left) and representative images of Dextran-Alexa555-trested iAstrocytes (right). Scale bar: 50μm.

**Figure S4.**
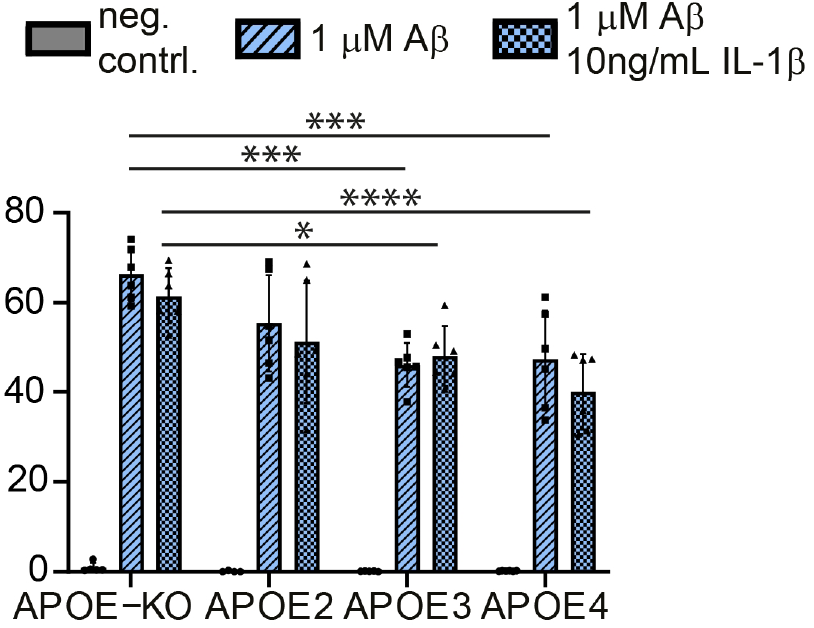
IL-1β activation does not affect allele-dependent Aβ uptake in iAstrocytes. Flow cytometry-based Aβ42 uptake assay showing percentage of Aβ positive cells. Cells were treated with 1μM pre-aggregated Aβ42-hilyte488 alone or pre-treated with 10ng/ml IL-1β. Data represent mean ± SD ***: p < 0.001; ****: p < 0.0001, Two-way ANOVA with post hoc Holm-Šídák’s multiple comparisons test).

**Table S1:** Statistical analysis of MSD. Table with results of statistical analysis and p-values for all comparisons in the MSD analysis of Figure 5D.

| | IFNγ | IL-10 | IL-12P70 | IL-13 | IL-1β | IL-2 | IL-4 | IL-6 | IL-8 | TNFα |
| --- | --- | --- | --- | --- | --- | --- | --- | --- | --- | --- |
| **NPC untreated** |  |  |  |  |  |  |  |  |  |  |
| APOE -/- vs. APOE 2/- | n.d. | n.d. | n.d. | n.d. | n.d. | 0.9718 | 0.9997 | 0.5948 | n.d. | n.d. |
| APOE -/- vs. APOE 3/- | n.d. | n.d. | n.d. | n.d. | n.d. | 0.9996 | >0.9999 | 0.9994 | n.d. | n.d. |
| APOE -/- vs. APOE 4/- | n.d. | n.d. | n.d. | n.d. | n.d. | 0.9994 | >0.9999 | 0.9879 | n.d. | n.d. |
| APOE 2/- vs. APOE 3/- | n.d. | n.d. | n.d. | n.d. | n.d. | 0.9875 | >0.9999 | 0.5222 | n.d. | n.d. |
| APOE 2/- vs. APOE 4/- | n.d. | n.d. | n.d. | n.d. | n.d. | 0.9893 | >0.9999 | 0.3966 | n.d. | n.d. |
| APOE 3/- vs. APOE 4/- | n.d. | n.d. | n.d. | n.d. | n.d. | >0.9999 | >0.9999 | 0.9967 | n.d. | n.d. |
| **Untreated** |  |  |  |  |  |  |  |  |  |  |
| APOE -/- vs. APOE 2/- | n.d. | n.d. | n.d. | n.d. | 0.9526 | 0.9956 | >0.9999 | 0.0283 | 0.9967 | >0.9999 |
| APOE -/- vs. APOE 3/- | n.d. | n.d. | n.d. | n.d. | 0.0458 | 0.9999 | >0.9999 | 0.1563 | 0.4446 | >0.9999 |
| APOE -/- vs. APOE 4/- | n.d. | n.d. | n.d. | n.d. | 0.9767 | 0.9976 | >0.9999 | <0.0001 | <0.0001 | 0.9997 |
| APOE 2/- vs. APOE 3/- | n.d. | n.d. | n.d. | n.d. | 0.1847 | 0.9984 | >0.9999 | 0.8815 | 0.5721 | >0.9999 |
| APOE 2/- vs. APOE 4/- | n.d. | n.d. | n.d. | n.d. | 0.999 | >0.9999 | >0.9999 | <0.0001 | 0.0002 | >0.9999 |
| APOE 3/- vs. APOE 4/- | n.d. | n.d. | n.d. | n.d. | 0.1096 | 0.9994 | >0.9999 | <0.0001 | 0.0081 | >0.9999 |
| **10ng/mL TNFα** |  |  |  |  |  |  |  |  |  |  |
| APOE -/- vs. APOE 2/- | <0.0001 | <0.0001 | 0.0148 | 0.0057 | 0.0001 | <0.0001 | 0.0011 | <0.0001 | 0.01 | n.d. |
| APOE -/- vs. APOE 3/- | <0.0001 | <0.0001 | 0.0075 | 0.8609 | <0.0001 | <0.0001 | <0.0001 | <0.0001 | <0.0001 | n.d. |
| APOE -/- vs. APOE 4/- | <0.0001 | <0.0001 | <0.0001 | <0.0001 | <0.0001 | <0.0001 | <0.0001 | <0.0001 | <0.0001 | n.d. |
| APOE 2/- vs. APOE 3/- | 0.4698 | 0.0203 | 0.9934 | 0.0629 | 0.0045 | 0.0004 | 0.4034 | 0.4084 | 0.0368 | n.d. |
| APOE 2/- vs. APOE 4/- | <0.0001 | <0.0001 | 0.0001 | <0.0001 | <0.0001 | <0.0001 | <0.0001 | <0.0001 | 0.0003 | n.d. |
| APOE 3/- vs. APOE 4/- | <0.0001 | <0.0001 | 0.0003 | <0.0001 | <0.0001 | <0.0001 | <0.0001 | <0.0001 | 0.3624 | n.d. |
| **10ng/mL IL1β** |  |  |  |  |  |  |  |  |  |  |
| APOE -/- vs. APOE 2/- | <0.0001 | <0.0001 | <0.0001 | 0.049 | n.d. | <0.0001 | <0.0001 | <0.0001 | >0.9999 | <0.0001 |
| APOE -/- vs. APOE 3/- | <0.0001 | <0.0001 | <0.0001 | 0.0013 | n.d. | <0.0001 | <0.0001 | <0.0001 | 0.0942 | <0.0001 |
| APOE -/- vs. APOE 4/- | <0.0001 | <0.0001 | <0.0001 | 0.0011 | n.d. | <0.0001 | <0.0001 | <0.0001 | 0.0501 | <0.0001 |
| APOE 2/- vs. APOE 3/- | 0.062 | <0.0001 | 0.0004 | 0.4914 | n.d. | <0.0001 | <0.0001 | <0.0001 | 0.1075 | <0.0001 |
| APOE 2/- vs. APOE 4/- | 0.0024 | <0.0001 | <0.0001 | 0.4495 | n.d. | <0.0001 | <0.0001 | <0.0001 | 0.0579 | <0.0001 |
| APOE 3/- vs. APOE 4/- | 0.5662 | 0.0006 | 0.036 | 0.9999 | n.d. | 0.0332 | <0.0001 | <0.0001 | 0.9924 | 0.9969 |

